# Length-dependent disassembly maintains four different flagellar lengths in *Giardia*

**DOI:** 10.1101/647115

**Authors:** SG McInally, J Kondev, Scott C. Dawson

## Abstract

How flagellar length regulation is achieved in multiciliated eukaryotic cells with flagella of different equilibrium lengths is unknown. The protist *Giardia lamblia* is an ideal model to evaluate length regulation as it has flagella of four different lengths. *Giardia* axonemes have both non-membrane-bound and membrane-bound regions, but lack transition zones. Here we quantified the contributions of intraflagellar transport (IFT)-mediated assembly and kinesin-13-mediated disassembly to length control. IFT particles assemble and inject at *Giardia’s* flagellar pore complexes, which act as diffusion barriers functionally analogous to the transition zone to compartmentalize the membrane-bound regions of flagella. IFT-mediated assembly is length-independent as train size, speed, and injection frequencies are similar between flagella of different lengths. In *Giardia*, kinesin-13 mediates a length-dependent disassembly mechanism of length regulation to balance length-independent IFT-mediated assembly, resulting in different lengths. We anticipate that similar control mechanisms are widespread in multiciliated cells where cytoplasmic precursor pools are not limiting.

## Introduction

Eukaryotic flagella and cilia (used interchangeably) are dynamic, membrane-bound and compartmentalized microtubule-based organelles that facilitate diverse cellular behaviors including motility, chemosensation, and directing hydrodynamic flow during development (Brooks & Wallingford, 2014; Pazour & Witman, 2003). Composed of over 500 distinct proteins, motile cilia are defined by the highly conserved axoneme structure consisting of nine microtubule doublets surrounding a central pair of microtubules (“9+2”) (Ishikawa, 2017). Axonemes are nucleated from mature centrioles or basal bodies, and extend from the complex transition zone (TZ), which acts diffusion barrier or ‘gate’ at the base of cilia (Reiter, Blacque, & Leroux, 2012). Due to its wide conservation in diverse microbial and multicellular flagellates, eukaryotic flagellar architecture likely predates the radiation of all extant lineages (Ishikawa, 2017; Sung & Leroux, 2013). Yet eukaryotic morphological diversity is vast, and many eukaryotes possess flagella with varied size, structure, and function presumably evolved from conserved flagellar components and assembly mechanisms (Avidor-Reiss, Ha, & Basiri, 2017; Ishikawa, 2017). Studies of flagellar assembly and length maintenance in diverse microbial eukaryotic lineages allow us to address a central question in evolutionary cell biology: how does cytoskeletal structural and functional variation arise from conserved cytoskeletal components and mechanisms?

Foundational studies in the green alga *Chlamydomonas reinhardtii* showed that axoneme assembly requires that the ciliary proteins synthesized in the cytoplasm are trafficked to the growing distal tip of the axoneme through a bidirectional, microtubule motor-driven process termed intraflagellar transport (IFT) (Kozminski, Johnson, Forscher, & Rosenbaum, 1993; Lechtreck, 2015). Assembly and maintenance of flagellar length is dependent upon IFT to provide building blocks to the site of assembly, the distal flagellar tip (Kozminski et al., 1993; Marshall & Rosenbaum, 1999). The components of IFT, including IFT particles, the BBsome, kinesin and dynein motors, and transition zone (TZ) complex proteins are widely conserved in free-living and parasitic unicellular flagellates such as *Tetrahymena*, *Leishmania*, *Trypanosoma*, and in the ciliated cell types of *C. elegans* and mammals (Buisson et al., 2013; Hao et al., 2009; Kozminski et al., 1993). Like variations in axoneme structure, variations in the IFT-mediated ciliogenesis mechanisms are also present in many flagellated unicellular protists—notably, such variations in axoneme number, structure, and assembly mechanism are hallmark features of many metazoan cell types (Brooks & Wallingford, 2014; Ishikawa, 2017). Atypical axoneme structure (e.g., primary cilia) can be the result of IFT-dependent mechanisms, and conversely the canonical axoneme (9+2) structure can be assembled through alternative IFT-independent “cytosolic ciliogenesis” mechanisms as described in apicomplexan parasites or in metazoan sperm (Avidor-Reiss & Leroux, 2015). Lastly, many common eukaryotes possess from two to thousands of cilia with diverse functions, lengths, morphologies, or inheritance patterns (e.g., the multiciliated protozoan *Tetrahymena* or multiciliated human epithelial cells). Is flagellar structural diversity explained simply by the fact that some lineages have tinkered with ancient, conserved IFT-mediated assembly and maintenance mechanisms, or have some flagellates evolved alternative flagellar assembly mechanisms?

The unicellular, parasitic protist *Giardia lamblia* is an ideal model organism to address the question of how unique flagellar types and flagellar lengths are built and maintained within a single multiciliated cell. *Giardia* has eight flagella organized as four bilaterally symmetric pairs with four different equilibrium lengths, implying a regulatory mechanism for sensing and differentially modulating assembly or disassembly rates. Equilibrium axoneme lengths of all eight flagella are also sensitive to microtubule stabilizing or depolymerizing drugs (Dawson et al., 2007). While each of the eight *Giardia* axonemes maintains the characteristic ‘9+2’ microtubule architecture, each axoneme also has a cytoplasmic, non-membrane-bound region that extends from a centrally located basal body before exiting the cell body as a membrane-bound flagellum (McInally & Dawson, 2016). Homologs of IFT–BBSome transport proteins, and kinesin-2 and dynein homologs are present in the genome, yet *Giardia* lacks a transition zone or TZ protein homologs (Avidor-Reiss & Leroux, 2015; Barker, Renzaglia, Fry, & Dawe, 2014). Lastly, during cell division and flagellar duplication, four intact mature axonemes and basal bodies are structurally inherited and four new axonemes are assembled *de novo* in each daughter cell (Hardin et al., 2017; Nohynková, Tumová, & Kulda, 2006). Cytoplasmic regions of the *de novo* posteriolateral and ventral axonemes are rapidly assembled prior to cytokinesis (Hardin et al., 2017).

The unique architecture and varied equilibrium lengths of *Giardia’s* eight flagella challenge canonical models of IFT-mediated flagellar assembly and length regulation. Flagella are classic models to study organelle size control, as each flagellum maintains a consistent steady-state, or equilibrium length, and the size of flagella can be represented in a single dimension, length (Marshall et al., 2005; Tamm, 1967). The prevailing explanation for regulation of equilibrium flagellar length is the “balance-point model”, which argues that constitutively controlled steady-state length is a balance between a length-dependent assembly rate and a length-independent disassembly rate (Marshall et al., 2005). Equilibrium length can be altered by modulating the rates of flagellar assembly or disassembly (Mohapatra, Goode, Jelenkovic, Phillips, & Kondev, 2016). The classic “long-zero” experiment in *Chlamydomonas* demonstrated length equalization of both flagella when one was amputated, defining the paradigm for flagellar length control across diverse eukaryotic cells (Coyne & Rosenbaum, 1970). Yet how are the four pairs of flagella in *Giardia* with *different* equilibrium cytoplasmic and membrane-bound flagellar lengths assembled and regulated in the same cell? Do multiciliated cells with different lengths require alternative mechanisms to assemble, maintain, and regulate distinct flagellar lengths?

To evaluate how *Giardia* differentially maintains and regulates different flagellar lengths, we quantified the dynamics of both IFT-mediated assembly and kinesin-13 mediated disassembly along the entire length of all axonemes – from the cytoplasmic basal bodies to the membrane-bound tips. By tracking IFT particle behavior and turnover in live cells in unprecedented detail, we discovered that the eight flagellar pore regions essentially act as the diffusion barriers for each flagellar compartment. IFT particles diffuse bidirectionally on the cytoplasmic regions and accumulate at each flagellar pore where IFT trains are assembled and injected into the membrane-bound axonemes, rather than at basal body or transition zone regions. We also determined that *Giardia* IFT train speed, size, and frequency of injection are similar regardless of the length of the flagellar pair. Increasing flagellar lengths with the MT-stabilizing drug Taxol also did not change the injection rate of IFT. *Giardia* kinesin-13 promotes microtubule disassembly and turnover of tubulin subunits at distal flagellar tips (Dawson et al., 2007), and here we show that kinesin-13 accumulates in a length-dependent manner to the eight flagellar tips. Thus *Giardia* maintains four different equilibrium flagellar lengths by tuning the rates of kinesin-13 mediated disassembly for specific flagellar pairs at the respective distal tips. Lastly, we propose a new model for flagellar length regulation that emphasizes the length-dependent disassembly process to balance a length-independent assembly rate for each flagellar pair. We anticipate that the innovations in flagellar length regulation in *Giardia* will echo known variations in flagellar structure, type, and number described in different human cell types and other flagellated microbial model systems.

## Results

### Four different equilibrium lengths of cytoplasmic and membrane-bound axonemes

The four flagellar pairs in *Giardia* have distinct lengths for both cytoplasmic and membrane-bound regions of axonemes. The cytoplasmic axonemal regions span from the basal body to flagellar pore (Figure 1A, shaded; 1B) and membrane-bound axonemal regions span from the flagellar pore to flagellar tip (Figure 1A, colored). To quantify the average lengths and the range of variation in lengths of both cytoplasmic and membrane-bound regions, we imaged populations of fixed trophozoites expressing a single, integrated copy of mNeonGreen-tagged β-tubulin to mark the MT cytoskeleton (Figure 1C). Total axoneme lengths (measured from basal body to flagellar tip) vary between the four pairs. The anterior axonemes had an average length of 19.1±0.4 µm, the caudal flagella were 20.5±0.6 µm, the posteriolateral flagella were 16.2±0.4 µm, and the ventral flagella were 16.8±0.9 µm (Supplemental Figure 1). We also confirmed the average lengths of membrane-bound regions in WBC6 trophozoites: Anterior flagella were 12.8±0.1 µm, caudal flagella were 8.7±0.2 µm, posteriolateral flagella were 8.1±0.1 µm, and ventral flagella were 13.7±0.9 µm (Figure 1D and (Hoeng et al., 2008)).

**Figure 1:**
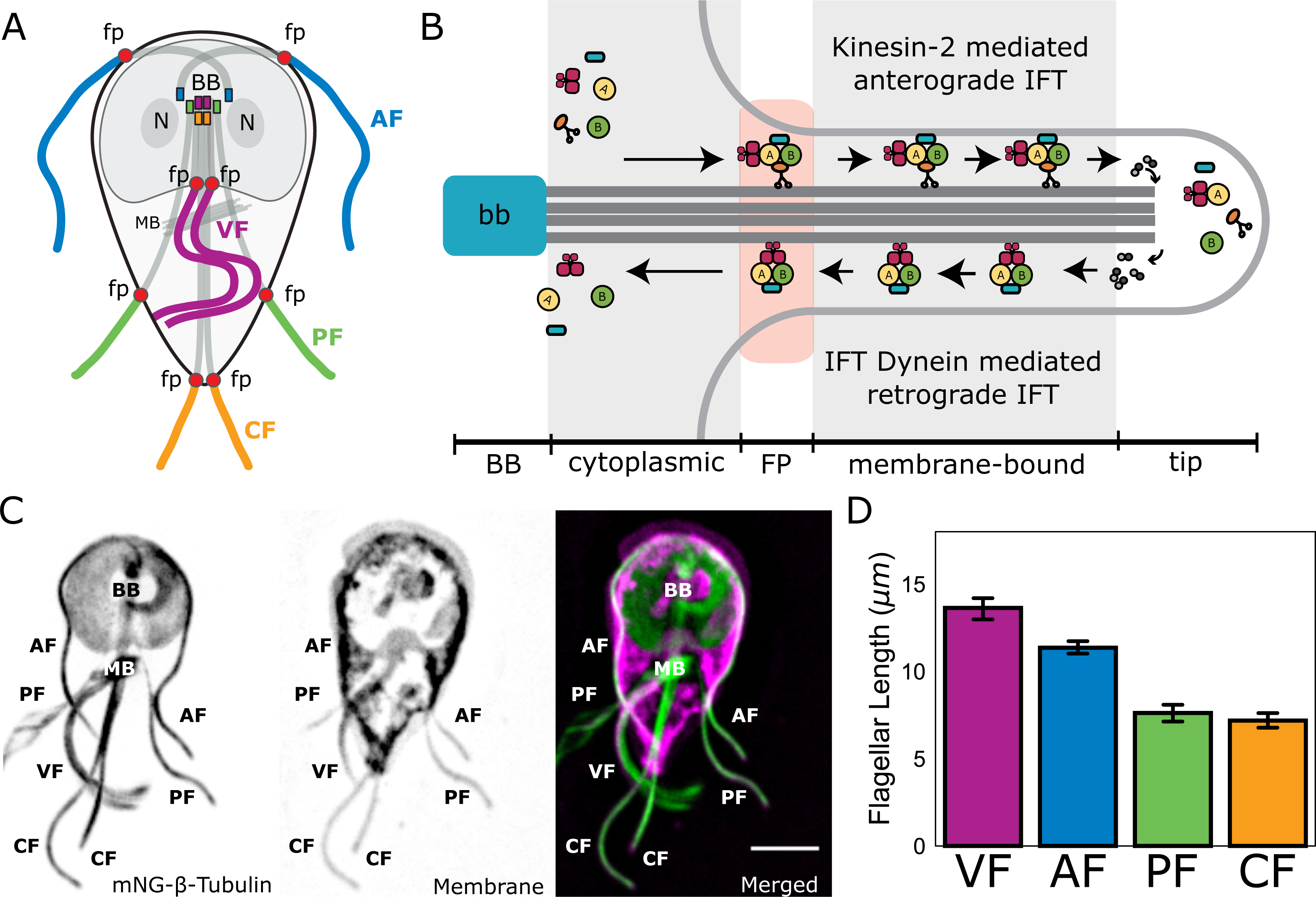
*Giardia* maintains four flagellar pairs at unique equilibrium lengths. (A) Schematic representation of membrane-bound, cytoplasmic, basal body (BB), and flagellar pore (fp) regions of the axoneme, as well as the two nuclei (N) and median body (MB). (B) Schematic representation of the specific regions of *Giardia*’s flagellar axoneme, including the cytoplasmic axoneme (cytoplasmic), flagellar pore (FP), membrane-bound axoneme (membrane-bound), and the flagellar tip (tip). (C) Fluorescent labeling of the microtubule cytoskeleton and membrane of a *Giardia lamblia* trophozoite, including the median body (MB), the basal body (BB), and the four flagellar pairs: anterior (AF), posteriolateral (PF), caudal (CF), and ventral (VF). Scale bar, 5µm. (D) Flagellar length quantification of membrane-bound regions of flagellar pairs of *Giardia* WBC6 trophozoites. The 95% confidence interval and average length are indicated. n ≥35 flagella for each pair. All pairs are statistically significantly different (p≤0.05, t-test) in membrane-bound length, except the posteriolateral and caudal flagella.

### All IFT homologs localize to both cytoplasmic and membrane-bound regions of the all flagella

IFT-mediated assembly of flagella requires the organization of IFT-A and IFT-B particles, kinesin and dynein motors, and the BBsome into multi-megadalton complexes called IFT trains (Figure 1B and (Lechtreck, 2015). While length maintenance of the membrane-bound regions of *Giardia’s* eight flagella is dependent on intraflagellar transport (IFT), the cytoplasmic regions are hypothesized to be assembled by an IFT-independent mechanism (Dawson et al., 2007; Hoeng et al., 2008). Previously, components of IFT-A (IFT140) and IFT-B (IFT81) were localized to both the cytoplasmic and membrane-bound axonemal regions (Hoeng et al., 2008). To confirm and extend this prior work, we imaged fluorescently tagged C-terminal fusions of all homologs of IFT particles (12), BBSome components (3), and kinesin-2 motor proteins (2). All *Giardia* IFT-A (IFT121, IFT122, IFT140) or IFT-B (IFT38, IFT54, IFT56, IFT57, IFT74/72, IFT80, IFT81, IFT88, and IFT172) particle homologs localized to the eight basal bodies, flagellar pores, flagellar tips, and along the lengths of both the cytoplasmic and membrane-bound regions of all axonemes (Figure 2A and (Hoeng et al., 2008)). Anterograde kinesin-2a and kinesin-2b motors localized to the eight flagellar pores and flagellar tips but had minimal localization to the cytoplasmic axonemes or basal bodies (Figure 2A and (Hoeng et al., 2008)). Furthermore, the localization puncta of IFT or kinesin-2 homologs at the flagellar tips are consistent with other studies of kinesin-2 distribution in the flagellum (Chien et al., 2017; Hendel, Thomson, & Marshall, 2018). Of the three BBSome homologs in *Giardia*, BBS4 localized primarily to the flagellar pores, BBS5 localizes primarily to the cytoplasmic axonemes, and BBS2 had cytoplasmic localization (Figure 2A). While distinct puncta of IFT trains were not observed on any of the cytoplasmic axonemes, all IFT proteins densely localized to cytoplasmic regions of all eight axonemes.

**Figure 2:**
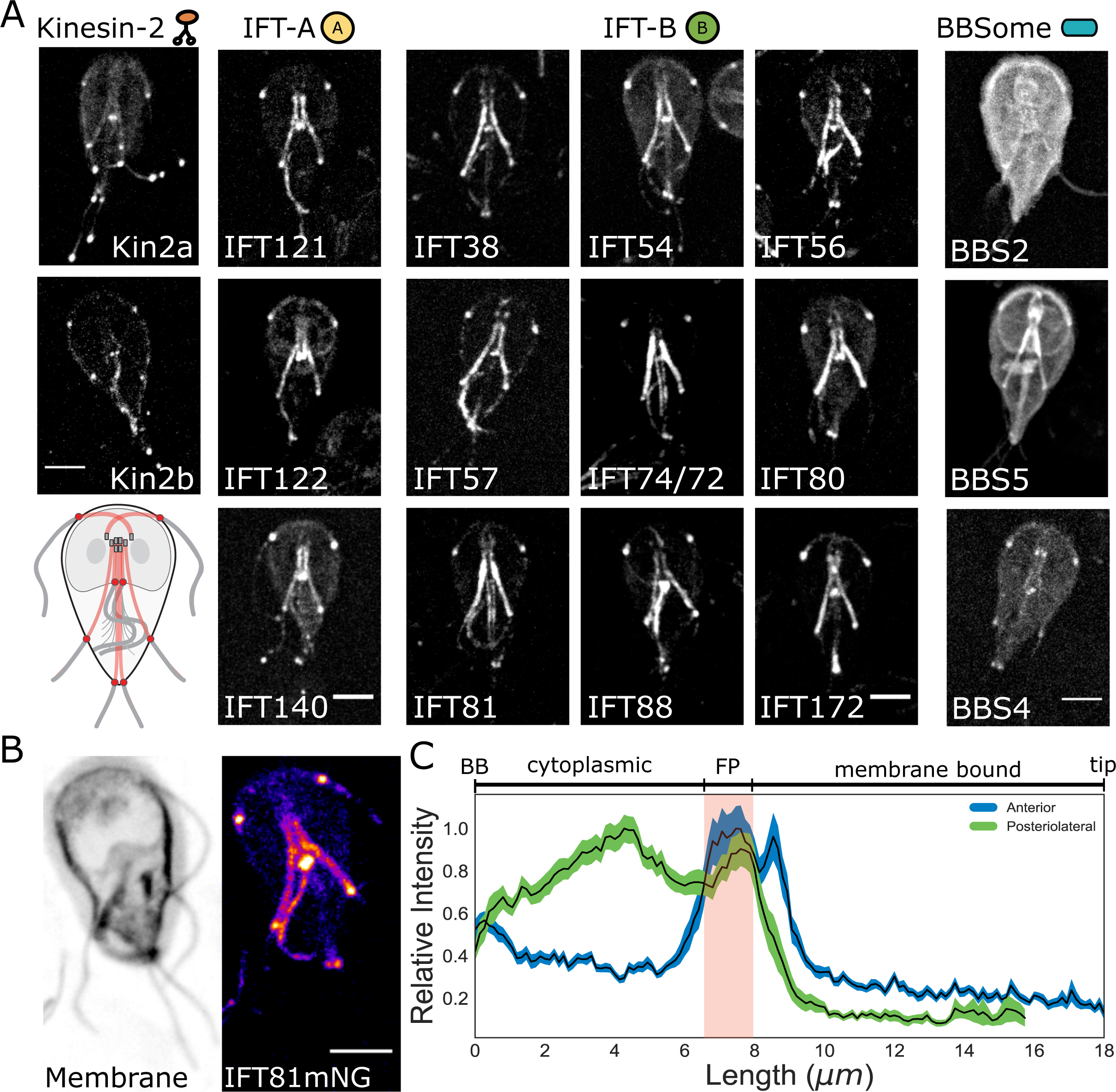
IFT particles accumulate at the flagellar pore in a flagellum-specific manner. (A) Maximum intensity projections of live cells show the distribution of kinesin-2, BBSome, IFT-A complex, and IFT-B complex proteins throughout the trophozoite. Representative schematic of IFT-A and IFT-B localizations is in the lowest left corner. All scale bars, 5µm. (B) IFT81mNeonGreen proteins are more concentrated at the flagellar pore regions of the flagellar pairs. Scale bar, 5µm. (C) Quantification of IFT81mNG distribution along the entire lengths of anterior and posteriolateral axonemes using line-scans. Black lines indicate mean intensity and shaded regions indicate 95% confidence intervals. Flagellar length is indicated on bottom axis and relative anatomical position is indicated on the top axis, with red shading indicating the flagellar pore region. n = 31 for each flagellar pair, from four independent experiments.

To determine whether overexpression of IFT proteins from episomal plasmids prevents observation of discrete IFT trains on the cytoplasmic axonemes, we also integrated mNeonGreen-tagged IFT81 (IFT81mNG) or GFP-tagged IFT81 (IFT81GFP) into the native *Giardia* IFT81 locus (Gourguechon & Cande, 2011) (Supplemental Figure 2). Strains expressing IFT81GFP or IFT81mNG from an integrated copy of the gene had the same subcellular localization as strains expressing tagged IFT81 from the episomal vector, but the labeling intensity was more uniform in the population of transformants. IFT81mNG was at least 3-fold brighter than IFT81GFP (Supplemental Figure 3).

### Axoneme specific accumulation of IFT particles at different flagellar pores

Due to the physical decoupling of the basal bodies and the membrane-bound regions of flagella in *Giardia*, the location of IFT train assembly and injection has remained unclear (Avidor-Reiss & Leroux, 2015). To determine the spatial distribution of IFT81mNG particles along the entire lengths of axonemes (basal body to flagellar tip), we traced fluorescence with line scans in two representative axonemal pairs (anterior and posteriolateral) in the IFT81mNG strain (Figure 2B). IFT81mNG fluorescence was not uniformly distributed along the lengths of either the anterior or posteriolateral axonemes (Figure 2B). The maximum fluorescence intensity for both flagellar pairs occurred at the flagellar pore, a region that lies at the transition from the cytoplasm to the compartmentalized flagellum (Figure 2C). IFT81mNG fluorescence in the anterior flagella had a single distinct maximum at the flagellar pore, whereas the posteriolateral flagella had one maximum at the posteriolateral flagellar pore and another maximum at a region immediately proximal to the ventral flagellar pores (Figure 2C). IFT81 localized less to the basal bodies of the anterior and posteriolateral axonemes than to the rest of the axoneme, in contrast to other flagellates (Hao et al., 2011; Prevo, Mangeol, Oswald, Scholey, & Peterman, 2015). The cytoplasmic regions of both the anterior and posteriolateral axonemes had higher intensities of IFT81mNG fluorescence than the membrane-bound regions, albeit less intensity than at flagellar pores. For all imaging experiments there was no difference in the localization of IFT81mNG fluorescence between either axoneme of each flagella pair. Our analyses of cytoplasmic axonemes were limited to the anterior and posteriolateral axonemes, as we were unable to reliably measure fluorescence intensity from the cytoplasmic regions of caudal and ventral flagella.

### The diffusive behavior of IFT particles in cytoplasmic regions results in IFT train assembly at each flagellar pore region

To understand how IFT particles accumulate in the flagellar pore regions in *Giardia*, we interrogated the behavior and turnover of IFT particles associated with the cytoplasmic axoneme and flagellar pore regions. Specifically, we used fluorescence recovery after photobleaching (FRAP) in the IFT81mNG strain to determine whether IFT particles on cytoplasmic axonemes are dynamic. After photobleaching the cytoplasmic regions of posteriolateral flagella (Figure 3A, 3C, Supplemental Video 1), we determined that IFT particles are dynamic on cytoplasmic axonemes and that recovery was also bi-directional. This analysis of turnover indicates that fluorescently labeled IFT81 molecules likely traverse the cytoplasmic axonemes in both anterograde and retrograde directions, despite our inability to detect distinct IFT trains traversing these structures using live imaging (Figure 3A). To assess the contribution of cytoplasmic axonemal IFT dynamics to the accumulation of IFT particles at each flagellar pore, we used FRAP to evaluate the turnover of IFT particles in the flagellar pores of posteriolateral axonemes (Figure 3B, 3D, Supplemental Video 2). IFT turnover was approximately three times faster at the flagellar pore (0.055±0.018 µm s^−1^) than in the cytoplasmic region (0.019±0.005 µm s^−1^) (Figure 3C, 3E).

**Figure 3:**
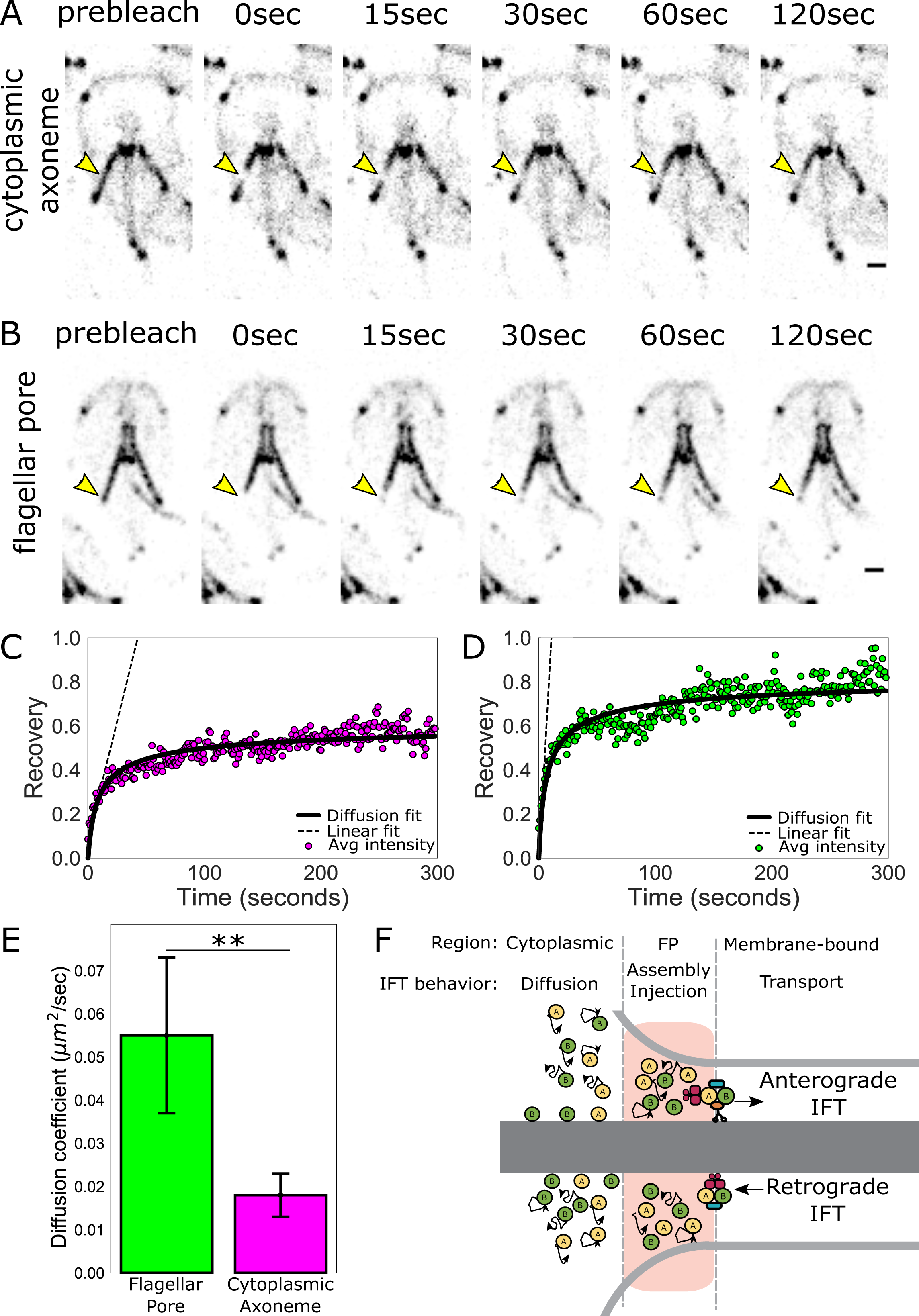
IFT train assembly occurs at the flagellar pore region. (A) Time series images of trophozoites expressing IFT81mNG prebleach, immediately post-bleach (0 sec, yellow arrow) of cytoplasmic axoneme or (B) flagellar pore regions, and during recovery (time in sec). Scale bar, 2µm. (C) Time averaged fluorescent recovery of posteriolateral cytoplasmic axonemes and (D) flagellar pores. Solid black lines indicate fit of the entire recovery phase. Dashed lines indicate linear fit of the initial recovery phase. n = 32 for bleached cytoplasmic axoneme regions, n = 25 for bleached flagellar pore regions, each from 3 independent experiments. (E) Estimated diffusion constants from fitting FRAP recovery of the flagellar pore and cytoplasmic regions of posteriolateral flagella. Means and 95% confidence intervals are indicated. Student’s t-test, **p<0.01. n≥25 cells, from ≥3 independent experiments. (F) Schematic representation of IFT particle behavior associated with the cytoplasmic axoneme, flagellar pore, and membrane-bound axoneme regions.

Differences in the rates of IFT particle turnover between the cytoplasmic and membrane-bound regions supports the idea that two processes promote the accumulation of IFT at the flagellar pore: (1) diffusion of IFT particles from the cytoplasmic axoneme region and (2) the return of IFT trains during retrograde transport (Figure 3F). To test this hypothesis, we developed a physical model that compares the relative contribution of these two processes to the accumulation of IFT particles at the flagellar pore. The model predicts a 3±1-fold difference in the initial linear-phase of the recovery between flagellar pore and cytoplasmic axoneme regions (Methods). We measured the slope of the initial linear-phase of recovery for these two regions and found 4.2±0.5-fold difference (Figure 3C, 3D). Therefore, our model provides additional support for the accumulation of IFT particles at flagellar pores due to the mixing of IFT particles diffusing from the cytoplasmic regions with IFT particles returning via retrograde transport from the membrane-bound axonemes (Figure 3F). The accumulation of IFT intensity combined with the absence of directed IFT transport from the cytoplasmic regions is also consistent with IFT train assembly occurring at each flagellar pore.

### IFT particle size, frequency and speed are similar between flagellar pairs of differing lengths

The balance point model of flagellar length control requires length-dependence of either the assembly rate, the disassembly rate, or both rates to establish an equilibrium length. Thus the maintenance of four different equilibrium flagellar lengths in *Giardia* could be due to either differential rates of IFT-mediated assembly or disassembly (Mohapatra et al., 2016). To evaluate length-dependent IFT-mediated assembly, we quantified and compared IFT dynamics within the membrane-bound compartment of three of the four flagellar pairs, each with different equilibrium lengths. We were unable to analyze IFT on the ventral flagella, which continued to beat while embedded (Figure 4A, Supplemental Video 3). Anterograde (Figure 4B, magenta) and retrograde (Figure 4B, green) IFT dynamics were compared using kymographs of the anterior, posteriolateral, and caudal flagella pairs (Figure 4B). For each flagellar pair, we quantified and compared the parameters of speed, size, and frequency of IFT trains undergoing transport within the membrane-bound flagellar compartment (Figure 4C-4F)(Engel, Ludington, & Marshall, 2009; Mangeol, Prevo, & Peterman, 2016).

**Figure 4:**
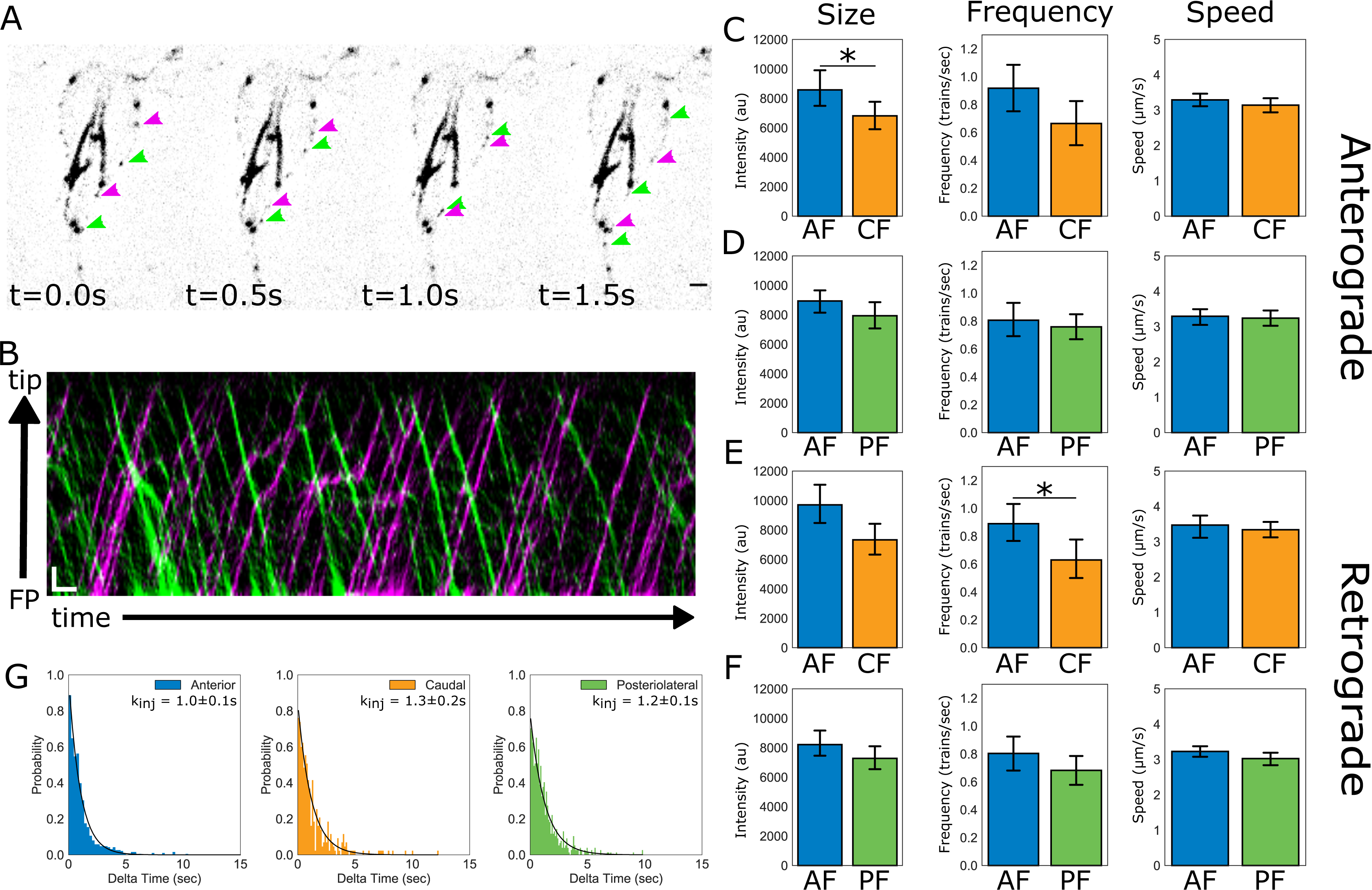
IFT dynamics are similar between flagellar pairs of different lengths. (A) Still images from time-lapse imaging of live trophozoites expressing IFT81mNG showing anterograde (magenta arrows) and retrograde IFT trains (green arrows). Scale bar, 2µm. (B) A representative kymograph of IFT train trajectories within the membrane-bound anterior flagellum. Total time is ∼26 sec. Scale bar, 1µm and 1 second. (C) Comparisons of anterograde IFT train intensity, frequency, and speed from anterior and caudal flagella. (D) Comparisons of anterograde IFT train intensity, frequency, and speed from anterior and posteriolateral flagella. (E) Comparisons of retrograde IFT train intensity, frequency, and speed from anterior and caudal flagella. (F) Comparisons of retrograde IFT train intensity, frequency, and speed from anterior and posteriolateral flagella. All plots show mean values with 95% confidence intervals. Student’s t-test, *p<0.05. n = 22 cells for the anterior and caudal flagella, n = 42 cells for the anterior and posteriolateral flagella, from N = 5 independent experiments. (G) Frequency histograms of the time-lag between IFT train injections for anterior (blue), posteriolateral (green), and caudal (orange) flagella. Black line indicates a fit to a single exponential equation to measure the injection rate for each flagellar pair. Injection rates are indicated with 95% confidence intervals.

For all flagella, the average IFT train speed was consistent for both anterograde and retrograde directions, and IFT train speeds were not different between pairs of different lengths. Specifically, the average anterograde IFT train speed for all flagellar pairs was between 3.0 – 3.2 µm/sec, with no significant differences between any of the measured flagella (Figure 4D). For all measured flagella, the retrograde IFT velocities were slightly greater than anterograde velocities, which is consistent with IFT velocities in other flagellates (Buisson et al., 2013). The average retrograde IFT velocity (∼3.2 µm/sec) was not significantly different between any of the measured flagellar pairs (Figure 4E).

In contrast, there were notable differences in the size of IFT trains between the flagellar pairs of different lengths, which could imply a length dependence of particle size for the anterior and caudal flagellar pairs. The average size of anterograde particles in anterior flagella was 24% larger than those in the shorter caudal flagella (Figure 4C). Retrograde IFT trains were also 21% larger in the anterior as compared to the caudal flagella (Figure 4E). Anterior anterograde or retrograde flagella IFT trains were also 11% larger as compared to the posteriolateral flagella (Figure 4D, F) yet this difference is not statistically significant.

Comparisons of anterograde IFT injection frequency between anterior, caudal, and posteriolateral flagella showed that IFT train injection is not significantly different between flagellar pairs of different lengths (Figure 4C, 4D). Retrograde IFT frequency was also not significantly different between flagellar pair of different lengths (Figure 4E, 4F). To compare and confirm IFT injection frequency rates (trains/sec) between different pairs, we also measured the time between each injection from kymographs filtered for only anterograde traffic (Figure 4B, magenta). The distribution of time-lag between injections is exponential, indicating a single rate limiting step for IFT train injection in the anterograde direction for the three pairs analyzed (Figure 4G). The frequency distribution was converted to a probability density function and a single exponential fit was used to measure the rate of injection (Methods, and Figure 4G). Overall, the frequency of anterograde IFT train injection was not significantly different between longer (anterior) and shorter (caudal, posteriolateral) flagellar pairs. The average time between IFT train injections was similar between the three flagellar pairs: 1.0±0.1 seconds for the anterior flagella; 1.3±0.2 seconds for caudal flagella; and 1.2±0.1 seconds for posteriolateral flagella (Figure 4G).

### Perturbation of flagellar length supports length independence of IFT injection

Next, we queried whether the total number of IFT trains within each flagellar pair scales linearly with length which is predicted by length-dependent IFT injection. For this analysis, we used 3D structured illumination microscopy (3D-SIM) to quantify the total integrated intensity of IFT trains in fixed trophozoites expressing integrated IFT81:GFP (Figure 5A) to provide an additional line of evidence for these observations. Fluorescence intensity of IFT81GFP and length of the membrane-bound regions of flagella were measured using line scans. The strong linear relationship between the total integrated fluorescence intensity and equilibrium flagellar length (R^2^=0.89) demonstrates that the total amount of IFT trains within each *Giardia* flagellum scale linearly with length (Figure 5B).

**Figure 5:**
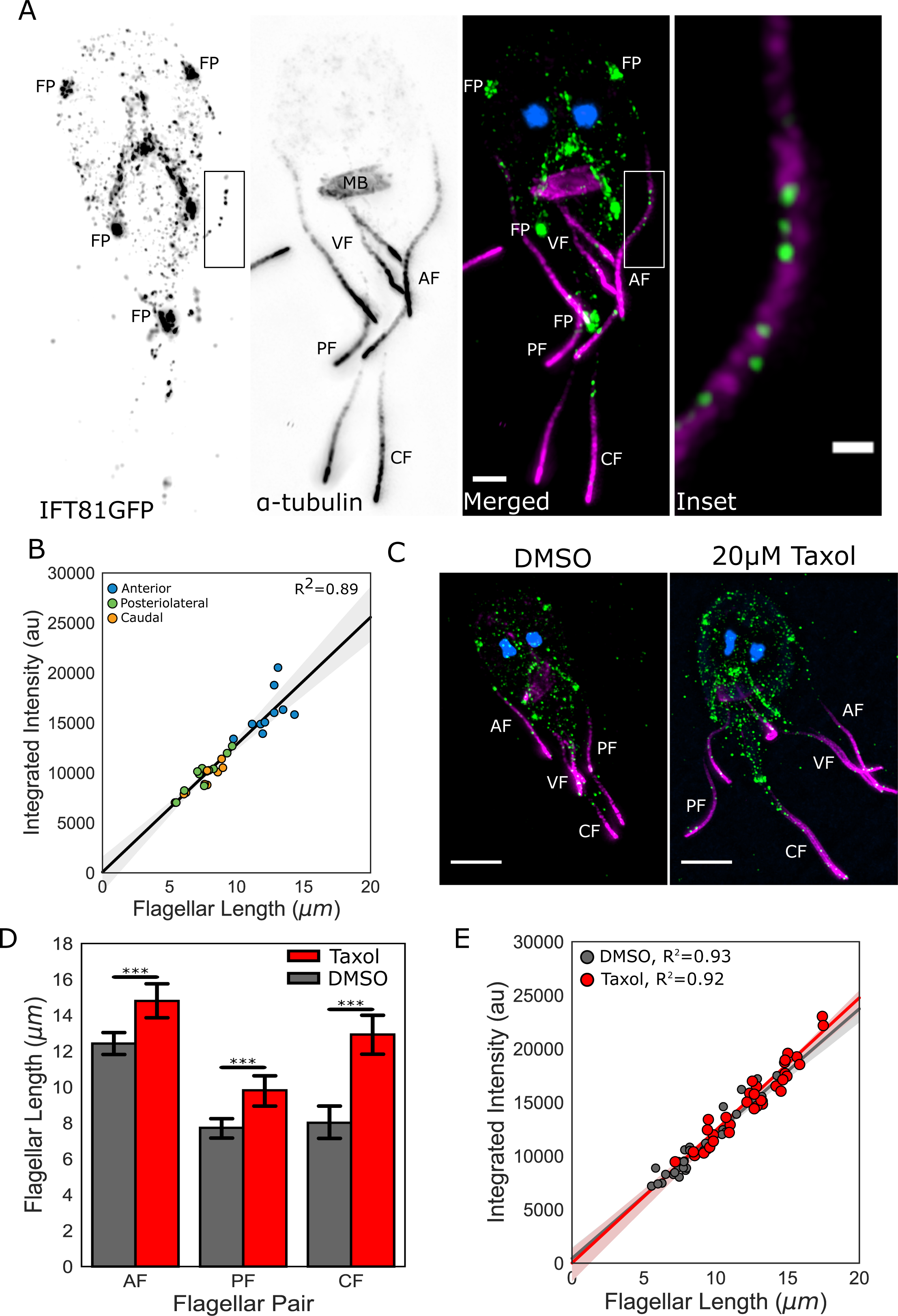
IFT injection is length-independent. (A) Representative structured illumination microscopy (SIM) image of IFT81GFP (green) immunostained for α-tubulin (magenta) and stained with DAPI (blue). Scale bar, 2µm. Boxed inset is enlarged on the right. Scale bar, 0.5 µm. (B) Total integrated intensity of IFT81GFP trophozoites plotted versus flagellar length. Orange dots indicate caudal flagella, green dots indicate posteriolateral flagella, and blue dots indicate anterior flagella. Linear fit (black line) and coefficient of determination are indicated. Shading indicates 95% confidence interval. (C) Representative SIM images of IFT81GFP trophozoites treated with DMSO (left) or 20µM Taxol (right) for 1 hour, then fixed and stained as in A. (D) Flagellar length of IFT81GFP trophozoites treated with DMSO (gray) or 20µM Taxol (red). Ten cells from three separate experiments were measured for each condition. Student’s t-test, ***p<0.001. (E) Total integrated intensity of IFT81GFP trophozoites treated with DMSO (gray) or 20µM Taxol (red) plotted versus flagellar length. Linear fit (gray and red lines) and coefficient of determination are indicated. Shading indicates 95% confidence interval.

*Giardia* axonemes are sensitive to microtubule assembly and disassembly dynamics during interphase. This offers another opportunity to evaluate the balance point model for length control in *Giardia*, because unlike other model systems (Wang et al., 2013), treatment with the microtubule-stabilizing drug Taxol rapidly increases all flagellar lengths (Dawson et al., 2007). Presumably, the pharmacological treatment with Taxol limits the frequency of catastrophe events at the dynamic plus-end of the microtubules in the axoneme (Dawson et al., 2007). Because we have shown that flagellar length scales with the total amount of IFT trains at equilibrium (Figure 5), we were able to test the response of the *Giardia*’s flagellar length control system to length perturbations in each flagellum induced by Taxol treatment (Figure 5 C-E). Treating trophozoites with 20 µM Taxol for one hour increased the average equilibrium flagellar length by 33% (Figure 5C, D). Specifically, the longer anterior flagella increased in length by 19%, the posteriolateral flagella by 27%, and shorter caudal flagella by 61% (Figure 5D). Following Taxol treatment, we again used line scans to quantify the total fluorescence intensity of IFT trains and the lengths of each flagellum (Figure 5C, 5E). The direct relationship between the total amount of IFT trains in each flagellum and its length was unchanged with Taxol treatment (Figure 5E), further supporting the length-independent rates of IFT injection in *Giardia*.

### Four different flagellar lengths are maintained by length-dependent disassembly

The balance point model of flagellar length control requires length dependence of either assembly and disassembly rates to achieve overall length control (Mohapatra et al., 2016). In *Chlamydomonas* and *Tetrahymena* length control is achieved through length-dependent IFT assembly with length-independent disassembly (Engel et al., 2009; Vasudevan et al., 2015). As we were unable to confirm length-dependent IFT-mediated assembly for the four *Giardia* flagellar pairs, we also investigated the potential contribution of length-dependent disassembly to the maintenance of four different equilibrium lengths. *Giardia* has a single kinesin-13 homolog that localizes to all distal flagellar tips, the median body, and the two mitotic spindles (Dawson et al., 2007). We have shown that kinesin-13 modulates flagellar disassembly in *Giardia*, as the overexpression of a dominant negative kinesin-13 rigor mutant and the depletion of kinesin-13 by CRISPRi-mediated knockdown cause increased flagellar lengths (Dawson et al., 2007; McInally et al., 2019).

To assess the contributions of kinesin-13 mediated disassembly to the maintenance of the lengths of all four flagellar pairs, we quantified all flagellar lengths in the CRISPRi-mediated kinesin-13 knockdown (K13kd) strain (Figure 6A, 6B). Using this K13kd strain, we have previously shown that caudal flagellar length increases (McInally et al., 2019). Here we extend this prior work to show that all flagellar pairs have steady-state length increases when kinesin-13 expression is inhibited by ∼60% (Figure 6A, 6B) (McInally et al., 2019), consistent with overexpression of a dominant negative rigor kinesin-13 mutant (Dawson et al., 2007). Compared to non-specific gRNA controls, the anterior flagella length in the K13kd strain is increased an average of 1.1±0.04 µm, the posteriolateral by 0.9±0.03 µm, the caudal by 3.1±0.18 µm, and the ventral by 1.6±0.06 µm (Figure 6H).

**Figure 6:**
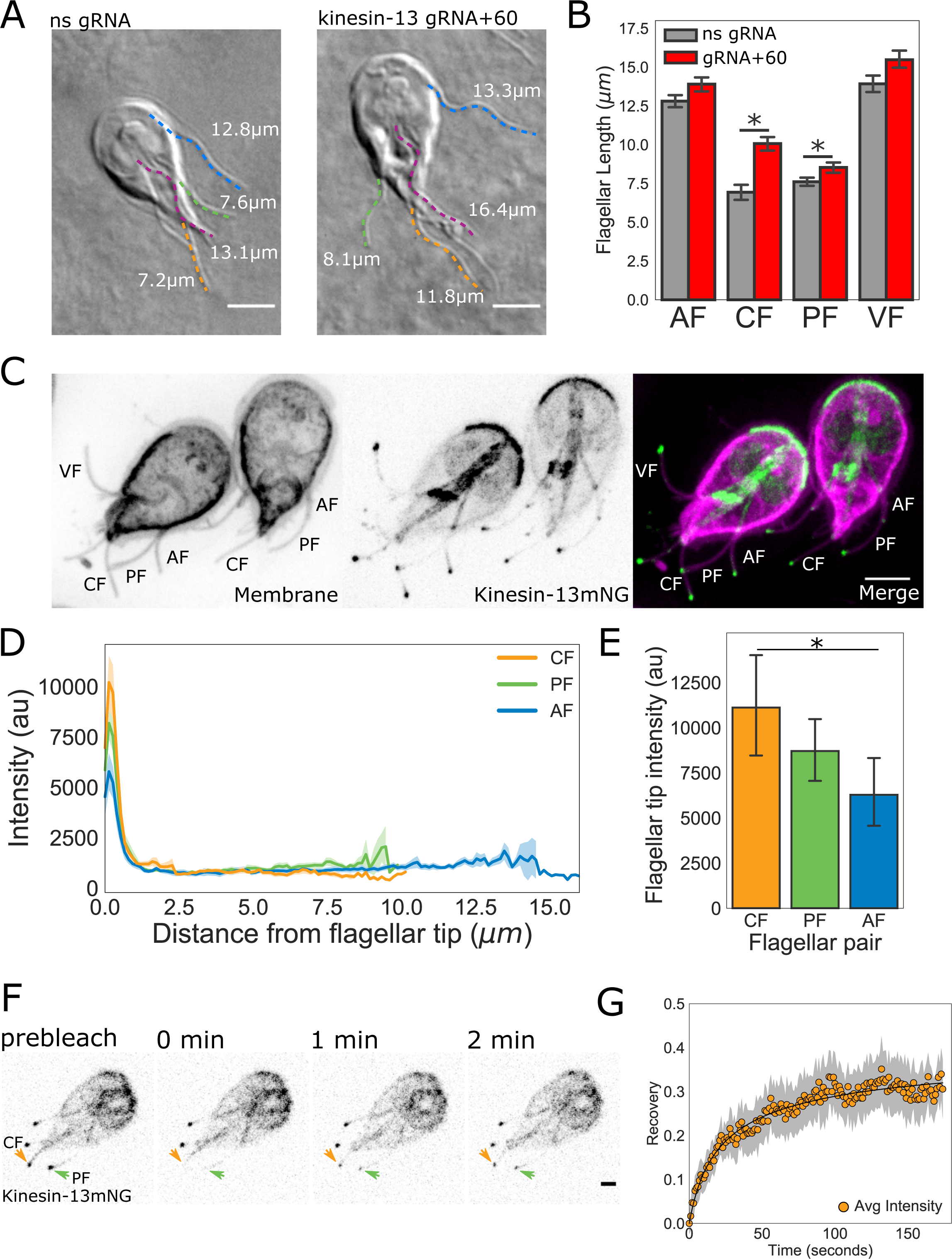
Kinesin-13 exponentially and dynamically localizes to the flagellar tip. (A) Representative images and quantification (B) of CRISPRi mediated knockdown of kinesin-13 (gRNA+60, red) as compared to a non-specific (ns, gray) gRNA. Blue traces indicate anterior flagella, magenta traces indicate the ventral flagella, green traces indicate the posteriolateral flagella, and orange traces indicate the caudal flagella. (C) Representative image of trophozoites expressing kinesin-13mNG with the cell membrane labeled to indicate the membrane-bound regions of the flagella. Scale bar, 5µm. (D) Kinesin-13mNG intensity profiles from the flagellar tip to the base of the membrane-bound regions of caudal (orange), posteriolateral (green), and anterior (blue) flagella. Shading indicates standard error of the mean. n≥23 for each flagellar pair, from two independent experiments. (E) Mean flagellar tip intensity plotted for each flagellar pair. 95% confidence intervals are indicated. Student’s t-test, *p<0.05. (F) Time series images of trophozoites expressing kinesin-13mNG prebleach, immediately post-bleach (0 sec, arrows) of caudal and posteriolateral flagellar tip regions, and during recovery (time in minutes). Scale bar, 2µm. (G) Time averaged fluorescent recovery of caudal flagellar tip regions following photobleaching. Solid black lines indicate fit of the entire recovery phase and shading indicates the 95% confidence interval. n = 19 caudal flagellar tips, from two independent experiments.

To quantify the localization and turnover dynamics of kinesin-13 along the lengths of all eight flagella, we constructed a fluorescent kinesin-13 fusion strain C-terminally tagged with mNeonGreen (kinesin-13mNG, Figure 6C). Kinesin-13mNG localized to all interphase microtubule structures, including the median body and ventral disc, and to both the cytoplasmic axonemes and the distal flagellar tips of all flagella (Figure 6C). To assess the contribution of kinesin-13-mediated disassembly to the maintenance of flagellar length in *Giardia*, we measured the spatial distribution of fluorescence within the membrane-bound region of the flagellum using line-scans from the flagellar tips to the flagellar pore. The maximum intensity of kinesin-13mNG fluorescence was at the distal region of the flagellar tips and intensity decreases exponentially within the first micrometer from the tip (Figure 6D). The shorter caudal flagella have more kinesin-13mNG at the distal tips than the longer anterior flagella (Figure 6E).

The specific localization of kinesin-13 at the distal flagellar tips suggests that this disassembly factor is actively transported to this region, likely via IFT. To characterize kinesin-13 transport dynamics, we used FRAP to determine the turnover of kinesin-13 at the distal tips of the caudal and posteriolateral flagella (Figure 6F, Supplemental Video 4). We observed fluorescence recovery of kinesin-13mNG at the distal flagellar tips within two minutes of photobleaching (Figure 6G). Coupled with the observed exponential decay of fluorescence signal from the distal flagellar tip, these findings support that kinesin-13 is transported to the distal flagellar tip.

### Distinguishing between two models of disassembly-driven flagellar length control

The dynamic localization of kinesin-13 at the flagellar tips coupled with a length-independent injection rate of IFT supports a length-dependent disassembly length control mechanism in *Giardia*. To further evaluate how a disassembly-driven mechanism involving the depolymerizing kinesin-13 could regulate four different equilibrium flagellar lengths, we developed two alternative models of length control. Both models assume that a higher disassembly rate is conferred by more kinesin-13 localized to shorter flagella (caudal, posteriolateral) than to longer (ventral, anterior) flagella during *de novo* assembly. Both models also propose that kinesin-13 and other cargo are transported to the flagellar tip via anterograde IFT as the flagellum elongates, but that kinesin-13 diffuses back to the flagellar base, independent of retrograde IFT (Figure 8A). Both assumptions are supported by the exponential distribution of kinesin-13 observed in the membrane-bound regions of the flagellum (Figure 6D) and turnover of kinesin-13 following photobleaching (Figure 6G).

The two models differ in how the assembly rate changes with the elongation of the flagella. The assembly rate is dependent on both the injection rate of IFT trains and the amount of structural precursor material, presumably tubulin, that remains in the cytoplasmic pool. The “Limited Precursor” model is motivated by length control studies in *Chlamydomonas* where there is a limiting pool of precursor material in the cytoplasm that is shared between the two flagella (Coyne & Rosenbaum, 1970; Marshall et al., 2005). As the flagella elongate the precursor pool is depleted, imparting an assembly rate that is length-dependent and decreases with flagellar length (Figure 7A, 7C). While we show that IFT injection is length-independent, the occupancy of IFT trains by structural components has been shown to vary, so that IFT trains may have cargo-binding sites that are unoccupied during transport (Wren et al., 2013). Therefore, length-independent injection of IFT trains does not necessarily indicate length-independent delivery of structural material to the distal flagellar tip. In the “Excess Precursor” model, there is an excess of precursor material; thus, the assembly rate is length-independent, and this rate remains constant during elongation (Figure 7B, 7D). As per the balance point model, it is the intersection of the assembly and disassembly rates that determine the steady-state length of flagella. Both proposed models achieve control of multiple flagella at distinct lengths (Figure 7C, 7D). However, the models differ in the expected concentrations of kinesin-13 localized to the flagellar tip at equilibrium and during elongation following Taxol treatment. We expect a greater concentration of kinesin-13 at the tips of short flagella in the “Limited Precursor” model, and that the concentration of kinesin-13 will decrease with Taxol elongation (Figure 7C). The “Excess Precursor” model predicts that the concentration of kinesin-13 will be similar between short and long flagella, and that these concentrations will not change with Taxol elongation of flagella (Figure 7D).

**Figure 7:**
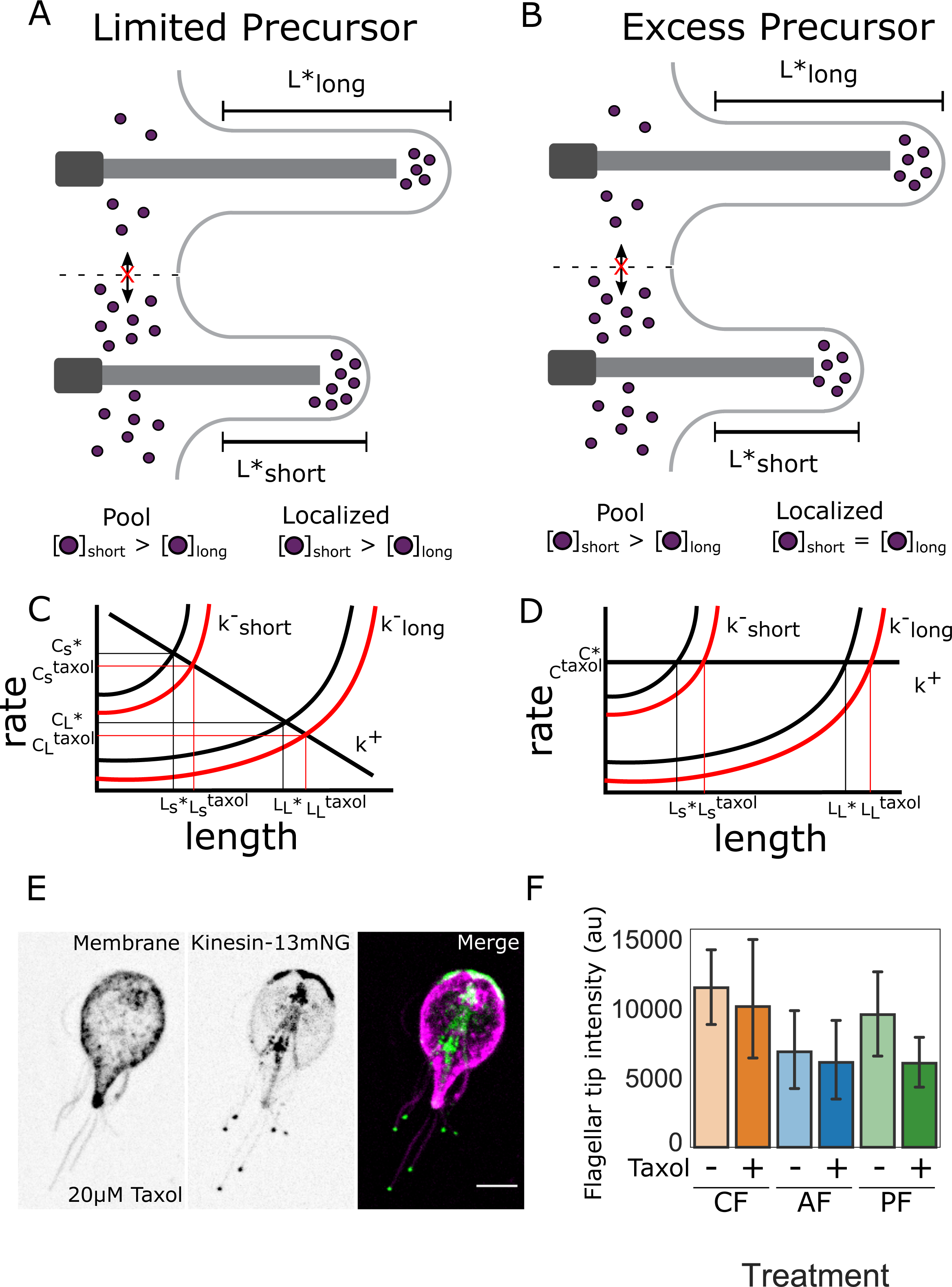
Length-dependent disassembly control flagellar length in *Giardia*. (A) Schematic representation of the “limited precursor” model. The pool of depolymerizing motors does not exchange between different flagella and the pool of depolymerizing motors is greater for shorter flagella than longer flagella at equilibrium. The concentration of depolymerizing motors is also greater at the tip of shorter flagella than longer flagella. (B) Schematic representation of the “excess precursor” model. The pool of depolymerizing motors does not exchange between different flagella and the pool of depolymerizing motors is greater for shorter flagella than longer flagella. The concentration of depolymerizing motors at the flagellar tip is equivalent for short and long flagella at equilibrium. (C, D) The rates of assembly (k^+^) and disassembly for short (k^−^_short_) and long (k^−^_long_) flagella as a function of length. (C) The assembly rate (k^+^) decreases as a function of length in the “limited precursor” model due to the depletion of the precursor from the pool as flagellar length increases. (D) The assembly rate (k^+^) is independent of length in the “excess precursor” model as this model assumes no depletion of the precursor pool. For both models the assembly and disassembly rates are balanced at distinct equilibrium lengths (L_s_^*^, L_L_^*^); the balance point also specifies the concentration of kinesin-13 at the tip (C*). Red lines indicate predicted changes to flagella length (L_s_^taxol^, L_L_^taxol^) and flagellar tip concentration of kinesin-13 (C_s_^taxol^, C_L_^taxol^) with Taxol induced elongation of flagella. (E) Representative image of trophozoites expressing kinesin-13mNG and treated with 20µM Taxol for 1 hour, with the cell membrane labeled to indicate the membrane-bound regions of the flagella. Scale bar, 5µm. (F) Flagellar tip intensity of kinesin-13mNG expressing trophozoites treated with DMSO (‘-‘) or 20µM Taxol (‘+’). n≥12 cells from two separate experiments were measured for each condition. Means and 95% confidence intervals are indicated.

To distinguish between these two models, we compared the fluorescence intensity of kinesin-13 at the flagellar tip during equilibrium and after flagellar elongation with Taxol (Figure 7E, 7F). The degree of flagellar length elongation with Taxol treatment was similar to prior measurements (Figure 5D, Supplemental Figure 4). We also plotted the fluorescence intensity of kinesin-13mNG versus the length of the flagellum (Supplemental Figure 4), and determined that there are no statistically significant differences between the fluorescence intensity of kinesin-13mNG at the flagellar tip at equilibrium or during Taxol elongation (Figure 7F), as predicted by the “Excess Precursor” model (Figure 7D). In the “Excess Precursor” model, the flagellum with more kinesin-13 at the base during *de novo* assembly balances the assembly rate at a shorter length than the flagellum with less kinesin-13 (Figure 7D).

## Discussion

Multiciliated *Giardia* trophozoites are a unique model in which to test how IFT-mediated assembly and disassembly mechanisms of flagellar length control enable different flagellar lengths in the same multiciliated cell. Like many model organisms, *Giardia* has canonical motile axonemes that are nucleated by basal bodies and have a conserved 9+2 axoneme structure. *Giardia* also possesses the majority of IFT, BBSome, and motor proteins (kinesin-2, kinesin-13, and IFT dynein) that are essential components of flagellar length control mechanisms in diverse model systems (Avidor-Reiss & Leroux, 2015; Lechtreck, 2015). Yet in contrast to other models, the eight *Giardia* axonemes are paired into four flagellar types with four different equilibrium lengths that include long, non-membrane-bound cytoplasmic regions. Flagellar beating of the different flagellar types is essential in *Giardia*’s life cycle for motility, cytokinesis, and excystation (Buchel, Gorenflot, Chochillon, Savel, & Gobert, 1987; Hardin et al., 2017; Lenaghan, Davis, Henson, Zhang, & Zhang, 2011). Each flagellar pair also has associated structural elements: the ventral flagella are associated with a fin-like density, the anterior and posteriolateral flagella have associated electron dense structures along their cytoplasmic regions, and the caudal complex surrounds the cytoplasmic regions of the caudal flagella (Elmendorf, Dawson, & McCaffery, 2003).

We demonstrate two types of evolutionary innovations in *Giardia* that may be broadly generalizable to other eukaryotes that generate variation in both axonemal architecture and in flagellar length regulation in multiciliated cells. First, while all *Giardia* flagella lack transition zone regions, each axoneme retains an analogous diffusion barrier defined by the eight flagellar pores to create a compartmentalized flagellum. This barrier has a similar function to the TZ but is not associated with basal bodies and is likely not homologous as the flagellar pore lacks TZ protein homologs. The second innovation enables four flagella of different lengths, and is a modification of flagellar length regulatory mechanisms, wherein kinesin-13 mediated disassembly rates are dependent on flagellar length, but IFT-mediated assembly rates are length-independent.

### Eight flagellar pores are cytoplasmic diffusion barriers analogous to the transition zone

The central cytoplasmic location of the eight basal bodies between the two nuclei is an atypical aspect of diplomonads like *Giardia*. In bi-flagellated *Chlamydomonas*, the basal bodies are anchored to the plasma membrane by transition zone fibers (Reiter et al., 2012). IFT particles (trains) are specialized for specific molecular cargos needed to build the growing axoneme. IFT trains accumulate in the transition zone, where they are loaded with axonemal structural material and transported to the distal flagellar tip by the heterotrimeric kinesin-2 complex (Deane, Cole, Seeley, Diener, & Rosenbaum, 2001; Wingfield et al., 2017). Retrograde trains are returned from the distal tip back to the cell body by cytoplasmic dynein (Pazour, Dickert, & Witman, 1999). Together, these motor-IFT complexes mediate dynamic trafficking of structural and signaling proteins into the compartmentalized flagellum, and are required for both flagellar assembly and maintenance (Lechtreck, 2015; Reiter et al., 2012). Regulation of IFT particle assembly, including regulation of motors and ultimately, flagellar length, is commonly attributed to regulatory proteins localizing to the transition zone. Thus the lack of a transition zone – known to both compartmentalize and concentrate proteins within the cilium – poses a conundrum in *Giardia*: how and where are IFT particles loaded and injected into the membrane-bound flagellar compartments (Avidor-Reiss & Leroux, 2015; Barker et al., 2014)?

Despite our prior analyses of flagellar assembly and disassembly in *Giardia* (Dawson et al., 2007; Hoeng et al., 2008), it has remained unclear where IFT particles assemble, mature, and are injected into the membrane-bound axonemal regions. Through GFP-tagging and imaging all IFT homologs in *Giardia*, we confirmed that all anterograde and retrograde IFT components localize not only to the membrane bound regions of axonemes, but also densely localize to the non-membrane-bound cytoplasmic regions (Figure 2). IFT particles also concentrate at each of the eight flagellar pores – the interface between the cytosol and compartmentalized membrane-bound axonemes (Figure 2A and (Hoeng et al., 2008)). Anterograde kinesin-2a and kinesin-2b motors also accumulate at each of the flagellar pores and flagellar tips, yet they lack a similar intensity of localization to the cytoplasmic regions of axonemes or basal bodies. The accumulation (Figure 2C) and turnover (Figure 3B, 3D) of IFT proteins at the flagellar pores result from the mixing of IFT particles from two sources: diffusive IFT particles from the cytoplasmic axonemal regions and IFT trains as they return via retrograde transport (Figure 3F). We propose that immature IFT particles first associate with cytoplasmic axonemal regions and mature at the eight flagellar pores, prior to their injection into the compartmentalized, membrane-bound axonemal region.

Thus, the eight flagellar pore regions act as a diffusion barrier between the cytoplasmic and membrane bound axoneme regions, resulting in the compartmentalization of the membrane-bound regions of the eight flagella. The mechanism by which the flagellar pores act as cytoplasmic diffusion barriers and regulate IFT injection is likely is analogous to that in other flagellates, yet in the absence of a TZ, *Giardia* must employ alternative or novel components. Overall, the TZ-independent accumulation and injection of IFT particles in *Giardia* demonstrates that eukaryotic flagella can be compartmentalized to concentrate proteins within the flagellar compartment in the absence of the canonical TZ complex, and thus raises questions regarding the necessity of this complex for IFT-mediated assembly in other flagellates. Further characterization of the ultrastructure and the composition of the flagellar pore complex in *Giardia* will be key to determining how this complex mediates compartmentalization and regulates IFT injection. The cytoplasmic face of pore complex also may play a role in directing the cytoplasmic axonemes to their defined exit points at the flagellar pores.

Other flagellated organisms lack IFT or TZ components and use IFT-independent or compartment-independent mechanisms to assemble flagella (Avidor-Reiss & Leroux, 2015), yet *Giardia’s* flagella appear to use a unique variation of these mechanisms. While an essential role for the TZ has been implied for many flagellates, there are several ciliated protozoans of the phylum *Apicomplexa* (e.g., *Plasmodium, Toxoplasma*) that also lack a transition zone structure. These apicomplexans, however, are thought to rely entirely on cytoplasmic ciliogenesis and do not possess compartmentalized flagella (Avidor-Reiss & Leroux, 2015). Other flagellated cells use both cytosolic and compartmentalized ciliogenesis, which creates two spatially distinct regions of the organelle that likely require unique assembly and maintenance dynamics. These observations support a mechanism of ciliogenesis that is not diffusion-limited and therefore does not require IFT for assembly of the cytoplasmic axonemal regions. Both mammalian and *Drosophila* sperm flagella employ variations of cytoplasmic ciliogenesis that require the invagination of the basal body into the cytoplasm following the initiation of compartmentalized ciliogenesis (Avidor-Reiss et al., 2017).

As in other eukaryotes with IFT-mediated flagellar assembly, conserved IFT components (e.g., IFT trains, BBsome, and kinesin and dynein motors) assemble the membrane-bound regions of each the eight *Giardia* axonemes (Hoeng et al., 2008; McInally et al., 2019). Cytoplasmic regions of each axoneme may be assembled in an IFT-independent manner as neither kinesin-2 knockdown nor expression of a dominant negative kinesin-2 affect cytoplasmic axoneme length (Dawson et al., 2007; Hoeng et al., 2008). Additionally, the cytoplasmic regions of the caudal and anterior axonemes of each daughter are proposed to be structurally inherited from the parental cell and are associated with mother, grandmother, and great-grandmother basal bodies (Nohynková et al., 2006). The two other flagellar pairs (ventral and posteriolateral) arise from *de novo* assembly immediately following cell division (Hardin et al., 2017; Sagolla, Dawson, Mancuso, & Cande, 2006). While we have not directly imaged the assembly of the cytoplasmic axonemal regions in this study, the lack of transport of IFT particles and kinesin-2 localization on cytoplasmic regions during interphase support an IFT-independent mechanism of flagellar assembly of the two *de novo* flagellar pairs (posteriolateral and ventral). This mode of ciliogenesis appears similar to that of *Plasmodium*, which occurs entirely in the cytoplasm with no membrane invagination or basal body migration in the absence of all known IFT components, including the transition zone and the BBSome (Avidor-Reiss & Leroux, 2015; Barker et al., 2014). The necessity of cytoplasmic ciliogenesis to precede the assembly of the membrane bound regions could essentially make the TZ dispensable in *Giardia*.

### Length-independent IFT-mediated assembly of all eight membrane-bound flagella

To set the length of a dynamic structure at a specific size, the rate of subunit addition must be balanced by the rate of subunit removal (Marshall et al., 2005; Mohapatra et al., 2016). The prevailing balance-point model of flagellar length control proposes that equilibrium length is achieved through the balance of assembly and disassembly rates (Engel et al., 2009; Ludington, Wemmer, Lechtreck, Witman, & Marshall, 2013; Mohapatra et al., 2016). The flagellar assembly rate is thus a result of IFT train injection into the membrane-bound region of the flagellum. IFT-mediated assembly is thought to be length-dependent as the size and number of IFT injections is inversely correlated with flagellar length (Engel et al., 2009). In contrast, the disassembly rate is thought to be length-independent (Engel et al., 2009; Ludington et al., 2013). Length-dependent assembly is proposed to arise from the depletion of the assembly motor, kinesin-2, at the flagellar base and the diffusive return of this essential IFT component from the flagella tip (Chien et al., 2017; Fai, Mohapatra, Kondev, & Amir, 2019; Hendel et al., 2018). In this way, the amount of kinesin-2 available to be incorporated into IFT trains acts as a length-ruler of flagella.

Again, the existence of eight flagella of four different equilibrium lengths (Figure 1) poses a challenge to the canonical model of flagellar length control. Based on the balance point model, one might predict that *Giardia* differentially regulates flagellar assembly amongst flagella of different lengths by differentially regulating IFT dynamics (particle size, number, or injection frequency). Yet we show here that IFT dynamics and IFT train injection are consistent between flagella of different equilibrium lengths (Figures 4, 5). Furthermore, this injection rate remains constant with increased flagellar length pharmacologically induced by Taxol treatment (Figure 5E). These observations imply that tuning of assembly rates is not a regulatory mechanism used to maintain differential lengths for different flagellar pairs in *Giardia*. The length-independent assembly of *Giardia* axonemes contrasts with observations in the green alga *Chlamydomonas reinhardtii*, wherein IFT train injection decreases with increasing flagellar length, therefore providing a length-dependent assembly rate (Chien et al., 2017; Hendel et al., 2018). Several kinases are known to regulate either assembly or disassembly rates in *Tetrahymena* and *Chlamydomonas* (CALK, LF4, Nrks/Neks), and *Giardia* has 198 Nek proteins whose functions are yet to be determined (Berman, Wilson, Haas, & Lefebvre, 2003; Bradley & Quarmby, 2005; Hilton, Gunawardane, Kim, Schwarz, & Quarmby, 2013; Manning et al., 2011; Meng & Pan, 2016; Wloga et al., 2006).

### Differential, length-dependent disassembly results in flagella of differing lengths

Kinesin-8 and kinesin-13 are depolymerizers of cytoplasmic, spindle, and ciliary microtubule ends (Helenius, Brouhard, Kalaidzidis, Diez, & Howard, 2006; Walczak, Gayek, Ohi, & Wordeman, 2013). Kinesin-13 depolymerizes the ends of microtubules via ATP hydrolysis and can traffic to microtubule ends by diffusion, by plus end–tracking proteins or by motors (Cooper & Schafer, 2000; Desai, Verma, Mitchison, & Walczak, 1999; Honnappa et al., 2009; Li et al., 2009). *Giardia* has a single kinesin-13 homolog that regulates dynamics of various MT arrays (e.g., two spindles, median body MTs, and axonemes). This is unlike metazoans, *Tetrahymena*, and trypanosomes, all of which have multiple kinesin-13 homologs with duplicated or differentiated functions in the various arrays. In *Giardia*, the ectopic expression of a dominant-negative, kinesin-13 rigor mutant or CRISPRi-mediated knockdown of kinesin-13 (Figure 6) results in dramatically longer flagella (Dawson et al., 2007; McInally et al., 2019). The ultrastructure of the all eight axonemes retain the canonical 9+2 arrangement of doublet microtubules but lack a flagellar tip complex (Dawson et al., 2007). Equilibrium flagellar lengths in *Giardia* are also sensitive to drugs that impact MT dynamics, and flagellar lengths are also increased by limiting MT catastrophes through treatment with the MT stabilizing drug Taxol (Dawson et al., 2007). Intrinsic microtubule dynamics may have a more pronounced contribution to flagellar length control in *Giardia* than in metazoans and *Chlamydomonas* as these flagella are comparatively less sensitive to microtubule stabilizing and destabilizing drugs (Wang et al., 2013).

Kinesin-13 contributions to flagellar assembly and disassembly have also been investigated in other microbial flagellates such as *Leishmania, Tetrahymena, and Chlamydomonas* (Blaineau et al., 2007; Chan & Ersfeld, 2010; Li et al., 2009; Vasudevan et al., 2015; Wang et al., 2013). Kinesin-13 has a similar role in mediating axoneme disassembly in *Leishmania*, where the knockdown of one (of the seven) kinesin-13 homologs promotes increased flagellar length, and overexpression of the same kinesin-13 homolog results decreased flagellar length (Blaineau et al., 2007). Depletion of the sole kinesin-13 in *Chlamydomonas* results in shorter flagella due to the depletion of the cytoplasmic tubulin pool required for IFT-mediated assembly, and also results in the disassembly of axonemes through IFT transport of kinesin-13 to the distal flagellar tip during the induction of flagellar resorption (Li et al., 2009; Wang et al., 2013). In *Tetrahymena* however, cell body MTs are shortened by kinesin-13, but this activity is not required for liberating ciliary precursor tubulin. Thus, in some flagellated cells, rather than directly playing a role in flagellar disassembly, kinesin-13 may indirectly impact flagellar IFT-mediated assembly, and ultimately flagellar length, through its role in the modulation of cytoplasmic tubulin precursor pools.

In *Giardia*, kinesin-13 also localizes to the median body, a centrally located cytoplasmic MT array, and to the kinetochore MTs of two mitotic spindles (Dawson et al., 2007). The median body is a semi-organized interphase MT array that is proposed to act as a cytoplasmic reservoir of tubulin prior to cell division (Hardin et al., 2017). Median body MTs are dynamic, and are sensitive to drugs that affect MT dynamics (Dawson et al., 2007). Over-expression of the kinesin-13 rigor mutant also results in decreases in median body volume (Dawson et al., 2007). In addition to regulating MT dynamics at the flagellar tips, kinesin-13 also regulates cytoplasmic tubulin pools through liberation of tubulin subunits in the median body in *Giardia*.

In the absence of differences in length-dependent axoneme assembly, we investigated the contribution of flagellar disassembly to flagellar length control in *Giardia*. Kinesin-13 dynamically localizes to the distal flagellar tip of all flagellar pairs, which is consistent with kinesin-13 depolymerizing in a length-dependent manner. In the absence of other active transport mechanisms within the membrane-bound flagellum, we propose that kinesin-13 is a cargo of IFT (Li et al., 2009). Due to turnover of kinesin-13 at the distal flagellar tip, coupled with the apparent exponential distribution of kinesin-13 fluorescence intensity within the flagellum, we suggest that kinesin-13 activity is a primary driver of differential flagellar length regulation in *Giardia* (Figure 8).

**Figure 8:**
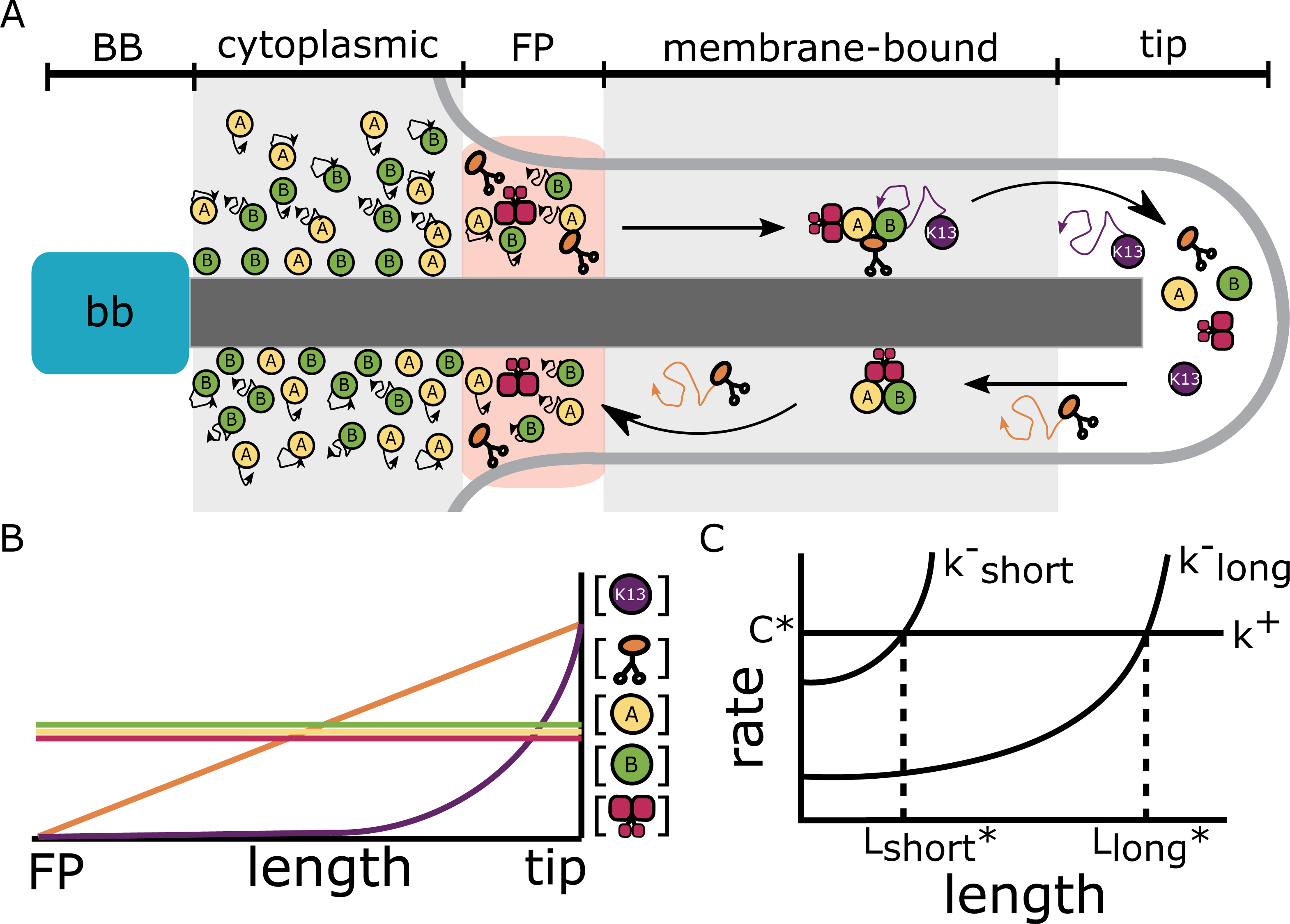
*Giardia* utilizes multimodal flagellar assembly and length-dependent disassembly to control four unique flagellar lengths. (A) Schematic of flagellar assembly and maintenance in *Giardia lamblia*. IFT particles move diffusively in the cytoplasmic axoneme regions. IFT trains are assembled in the flagellar pore region and are injected into the membrane-bound region of the axoneme. Within the membrane-bound region, IFT particles undergo anterograde transport via kinesin-2 mediated transport until they reach the distal flagellar tip. IFT trains are reorganized into retrograde directed trains and carried back to the flagellar base by IFT dynein. Kinesin-2 and kinesin-13 are not included in retrograde IFT trains, and instead diffuse back to the flagellar base. While kinesin-2 can freely diffuse to the flagellar base, kinesin-13 can be ‘recaptured’ by anterograde IFT trains and carried back to the distal tip (Naoz et al., 2008). (B) Predicted fluorescence intensity profiles of the various IFT components in the membrane-bound region of the flagellar axoneme. Bi-directional transport gives a fluorescent signal that does not change with length (IFT-A, IFT-B, IFT dynein). Uni-directional transport coupled with free diffusion is expected to give a profile that decreases linearly from the flagellar tip to the base (Kinesin-2). Uni-directional transport coupled with diffusion and anterograde recapture gives a profile that decreases exponentially from the distal flagellar tip (Kinesin-13) (Naoz et al., 2008). (C) The rates of polymerization (k^+^) and depolymerization for short (k^−^_short_) and long (k^−^_long_) flagella as a function of length. The polymerization rate is length-independent, while depolymerization rates are length-dependent. These rates intersect to give two distinct equilibrium lengths (L_short_^*^, L_long_^*^) with equal concentrations of kinesin-13 at the distal flagellar tip.

### A disassembly mediated model for flagellar length control

*Giardia* tunes differential flagellar lengths through the modulation of axonemal-specific, length-dependent flagellar disassembly rates, rather than length-dependent IFT-mediated assembly rates as reported in other systems (Figure 8C). The four equilibrium flagellar lengths are achieved by modulating the amount of kinesin-13 accumulated to the distal flagellar tip during assembly (Figure 8C); however, it remains unclear how kinesin-13 is differentially transported or regulated at the eight different flagellar tips.

Generally, we propose that kinesin-13 is transported to the distal flagellar tips by IFT, where it is released from the IFT train and can interact with the flagellar axoneme to promote disassembly. When unbound from the axoneme, kinesin-13 diffuses back toward the flagellar pores, similar to kinesin-2 (Chien et al., 2017; Hendel et al., 2018). During this diffusion, kinesin-13 can rebind to an anterograde IFT train, preventing the “escape” of kinesin-13 from the distal flagellar tip and generating the observed distribution and tip accumulation within the flagellar compartment (Figure 8A). Importantly, this model of flagellar length assembly and maintenance generates specific predictions about the intensity profiles of the various IFT components within the membrane-bound region of the flagellum. Components that undergo anterograde and retrograde transport (IFT-A, IFT-B, IFT dynein) are expected to display a uniform fluorescence distribution within the membrane-bound flagellum (Figure 8B). Anterograde transport coupled with retrograde diffusion produces a linear decrease is fluorescence signal intensity and would be expected for kinesin-2 (Figure 8B, (Chien et al., 2017; Hendel et al., 2018)). Lastly, we propose that components which undergo anterograde transport with retrograde diffusion and recapture by anterograde complexes are expected to display an exponential decrease in fluorescence signal (Naoz, Manor, Sakaguchi, Kachar, & Gov, 2008), as is observed with kinesin-13 (Figure 8B).

Both between species and even within cell types of the same species, eukaryotic cells have diverse cytoskeletal architectures that enable innovations in motility and other cellular functions. Defining the molecular mechanisms by which non-canonical flagellated cells like *Giardia* alter the balance of well-studied IFT assembly and kinesin-13 mediated disassembly mechanisms illuminates how cells can evolve varied morphological forms. Beyond microbial flagellates, we expect the mechanistic and structural innovations we describe in *Giardia* will echo well-described variations in flagellar structure, type, and number found in different cell types in humans and other multicellular model systems.

## Methods

### Strains and culture conditions

*Giardia lamblia* (ATCC 50803) strains were cultured in modified TYI-S-33 medium supplemented with bovine bile and 5% adult and 5% fetal bovine serum [56] in sterile 16 ml screw-capped disposable tubes (BD Falcon). Cultures were incubated upright at 37°C without shaking as previously described(Hagen et al., 2011). GFP and mNeonGreen-tagged IFT strains were created by electroporation of episomal vectors into strain WBC6 using approximately 20 µg plasmid DNA (Hagen et al., 2011). Tagged strains were maintained with antibiotic selection (50 µg/ml puromycin and/or 600 µg/ml G418)(Hagen et al., 2011). Trophozoites treated with Taxol were grown to confluency and split into 6 ml culture tubes 2 hours prior to incubation with Taxol (20µM final concentration) or DMSO (0.2% final concentration) for one hour. Live or fixed imaging was performed as described below.

### Construction of episomal and integrated C-terminal GFP and mNeonGreen-tagged IFT strains

Eleven intraflagellar transport (IFT) homologs were identified by homology searches in the *Giardia* genome (GL50803) using GiardiaDB (Aurrecoechea et al., 2009). C-terminal GFP tagged IFT homologs were created by PCR amplification of genomic DNA (Table 2) and subsequent cloning of the IFT homolog amplicons into a *Giardia* Gateway cloning vector (Hagen et al., 2011). For live imaging of IFT particles, we constructed an IFT81mNeonGreen strain by PCR amplification of the IFT81 gene (GL50803_15428) (Table2) to generate IFT81 flanked by 5’ NotI and 3’ BamHI restriction sites. The resulting amplicon was gel purified using a Zymoclean Gel DNA Recovery kit (Zymo Research) and cloned into pKS_mNeonGreen-N11_PAC (Hardin et al., 2017; Shaner et al., 2013) using Gibson cloning (Gibson et al., 2009). The resulting IFT81mNeonGreen and IFT81GFP plasmids were linearized with AflII for integration into the native locus (Gourguechon & Cande, 2011). To verify integration, total genomic DNA was extracted from tagged IFT81 strains using DNA STAT-60 (Tel-Test, Inc.), and integration of the C-terminal tag was confirmed by PCR amplification (Supplemental Figure 2).

### Immunostaining and light microscopy

*Giardia* trophozoites were grown to confluency as described above. Media was then replaced with 1x HBS (37°C), and the cultures were incubated at 37°C for 30 minutes. To detach and harvest cells, culture tubes were incubated on ice for 15 minutes and centrifuged at 900 × g, 4°C for five minutes. Pellets were washed twice with 6 ml cold 1x HBS and resuspended in 500 µl 1x HBS. Cells (250 µl) were attached to warm coverslips (37°C, 20 min), fixed in 4% paraformaldehyde, pH 7.4 (37°C, 15 min), washed three times with 2 ml PEM, pH 6.9 (Woessner & Dawson, 2012) and incubated in 0.125M glycine (15 min, 25°C) to quench background fluorescence. Coverslips were washed three more times with PEM and permeabilized with 0.1% Triton X-100 for 10 minutes. After three additional PEM washes, coverslips were blocked in 2 ml PEMBALG (Woessner & Dawson, 2012) for 30 minutes and incubated overnight at 4°C with anti-TAT1 (1:250) and/or anti-GFP (1:500, Sigma) antibodies. The following day, coverslips were washed three times in PEMBALG and then incubated with Alexa Fluor 555 goat anti-rabbit IgG (1:1000; Life Technologies), Alex Fluor 594 goat anti-mouse antibodies (1:250; Life Technologies) and/or Alex Fluor 647 goat anti-mouse (1:250; Life Technologies) antibodies for 2 hours at room temperature. Coverslips were washed three times each with PEMBALG and PEM and mounted in Prolong Gold antifade reagent with DAPI (Life Technologies).

### Flagellar pair length measurements

*Giardia* trophozoites were fixed and stained with TAT1 (1:250) and Alexa Fluor 594 goat anti-mouse IgG (1:250; Life Technologies) as described above. For flagellar pair length measurements, serial sections of immunostained trophozoites were acquired at 0.2 µm intervals using a Leica DMI 6000 wide-field inverted fluorescence microscope with a PlanApo ×100, 1.40 numerical aperture (NA) oil-immersion objective. DIC images were analyzed in FIJI (Schindelin et al., 2012) using a spline-fit line to trace the flagella from the cell body to the flagellar tip. Flagellar length measurements were analyzed and quantified using custom Python scripts. n ≥35 flagella for each pair. Flagellar length data are shown as mean relative length changes with 95% confidence intervals.

### Live imaging of IFT in Giardia

For live imaging, strains were grown to confluency, incubated on ice for 15 minutes to detach cells and pelleted at 900 × g for five minutes at 4°C. Cell pellets were washed three times in 6 mL of cold 1x HBS. After the final wash, cells were resuspended in 1mL of cold 1x HBS. For live imaging, 500µL of washed trophozoites were added to the center of a prewarmed 35 mm imaging dish (MatTek Corporation) and incubated for 20 minutes at 37°C. The imaging dish was washed with three times with warmed 1x HBS to remove unattached cells. For some experiments, CellMask Deep Red plasma membrane stain (ThermoFisher) was used to label the cell membrane (1x final concentration, 15 minutes at 37°C). Attached cells were embedded in 1mL 3% low melt agarose (USB Corporation) in 1x HBS (37°C) to limit flagellar beating and prevent detachment. The imaging dish was sealed using parafilm and imaged on a 3i spinning disc confocal microscope (Intelligent Imaging Innovations, Inc.).

### Quantification of IFT81mNG full axoneme fluorescence intensity

For analysis of IFT along the entire length of axonemes (basal body to flagellar tip), the IFT81mNG strain was grown to confluency and prepared for live imaging (see above). Images were acquired on a 3i spinning disc confocal microscope as described below. The segmented line tool in FIJI was used to measure the fluorescence intensity along the entire axoneme from basal body to the flagellar tip. Quantification was conducted for the anterior and posteriolateral flagella from 31 cells acquired in three independent experiments. The overall distance from the basal body to the flagellar pore and the flagellar tip was also recorded for each cell. Python scripts were used to plot the mean intensity for all analyzed cells and to calculate and plot the 95% confidence interval for all measurements.

### Fluorescence recovery after photobleaching (FRAP)

To quantify IFT dynamics at different regions of axonemes, the integrated IFT81mNeonGreen strain was grown to confluency and prepared for live imaging as described above. Images were acquired using a 3i spinning disc confocal microscope (Intelligent Imaging Innovations, Inc.) using a 63x, 1.3 NA objective. The microscope was warmed to 37°C one hour prior to image acquisition to maintain cells at physiological temperature and DefiniteFocus (Zeiss) was used to prevent drift during image acquisition. Either the cytoplasmic axoneme region or the flagellar pore region of the posteriolateral flagellum was bleached using Vector (Intelligent Imaging Innovations, Inc.) with a 488nm laser (10% laser power, 5ms exposure). To monitor recovery, images were acquired with a 488nm laser (50ms exposure and 50% laser power on gain set to 2 with intensification set to 667) at one-second intervals. Images were processed using the Template Matching plugin for FIJI to correct drift during acquisition. Once corrected for drift, ROIs were identified, and intensity measurements were recorded for each experimental time point. Background intensity measurements were taken for each cell analyzed from an adjacent area with no detectable fluorescence. Photobleaching measurements were taken from a non-bleached region of the cell analyzed. Quantification of fluorescence recovery was performed using custom Python scripts to subtract background intensity and correct for photo-bleaching. Individual recoveries were fit using Equation 1 (Ellenberg et al., 1997) to measure the diffusion constant:

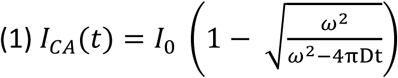

where *I_CA_*(*t*) is the intensity as a function of time and zero time is bleaching event; *I*_0_ is the final intensity after recovery; ω is the bleach strip width; D is the diffusion constant.

At short times, Equation 1 becomes Equation 2:

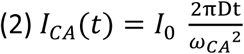

where ω_CA_ is the bleach strip width for the cytoplasmic axoneme. To predict the initial recovery of the flagellar pore region, the flux from retrograde IFT transport is added to Equation 2:

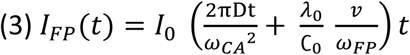

where *I_FP_*(*t*) is the intensity as a function of time; 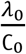 is the relative difference in integrated fluorescence intensity in the membrane-bound axoneme and the cytoplasmic axoneme; *v* is the speed of retrograde IFT; and ω_FP_ is the width of the bleach strip for the flagellar pore. Inputting all measured parameters into Eqs 2 and 3 predicts that 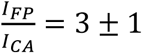 for the initial period of recovery. Fits of the initial linear phase of recovery were conducted using linear regression.

All recovery measurements (32 cytoplasmic axonemes and 25 flagellar pores, from at least 3 independent experiments) were average and rescaled based on the average initial intensity and bleach depth.

### IFT particle tracking using kymograph analysis

To track IFT train traffic in live cells, the integrated IFT81mNeonGreen strain was first grown to confluency and then prepared for live imaging as described above. Images were acquired using a 3i spinning disc confocal microscope (Intelligent Imaging Innovations, Inc.) using a 100x, 1.46 NA objective (330 images, 30ms exposure, intensification = 667, and gain = 2; the total time was about 26 seconds for each acquisition for a frame rate of approximately 13fps). The microscope was warmed to 37°C one hour prior to image acquisition to maintain cells at physiological temperature and DefiniteFocus (Zeiss) was used to prevent drift during image acquisition.

Kymographs were generated using the KymographClear 2.0 plugin for FIJI (Mangeol et al., 2016). A maximum intensity projection image was generated from the time lapse series to identify flagella, and a segmented, spline-fit line was used to trace identified flagella in the time-lapse images. Kymographs were generated for cells that had at least two identified flagella from different flagellar pairs to make intracellular comparisons. Intracellular comparisons were made for 22 cells for the anterior and caudal flagella and for 42 cells for the anterior and posteriolateral flagella, obtained from five independent microscopy experiments. Kymographs were analyzed using KymographDirect software with background correction and correction for photobleaching, and data were further analyzed using custom Python scripts.

### IFT injection frequency distributions

Forward-filtered kymographs generated above were used to measure the time-lag between IFT train injections. Using custom Python scripts, we calculated the time between each injection event for anterior, posteriolateral, and caudal flagella. Frequency histograms for each flagellar pair were fit with a single exponential to measure the rate of IFT injection for each flagellar pair.

### Quantification of integrated IFT particle intensity using super-resolution microscopy

The integrated IFT81GFP strain was first grown to confluency, then fixed and stained as described. 3D stacks were collected at 0.125µm intervals on a Nikon N-SIM Structured Illumination Super-resolution Microscope with a 100x, 1.49 NA objective, 100 EX V-R diffraction grating, and an Andor iXon3 DU-897E EMCCD. Images were reconstructed in the “Reconstruct Slice” mode and were only used if the reconstruction score was 8. Raw and reconstructed image quality were further assessed using SIMcheck and only images with adequate scores were used for analysis (Ball et al., 2015).

To determine intensity profiles along the length of flagellar pairs, we used the maximum intensity projections of reconstructed SIM images for tubulin (anti-TAT) and IFT81GFP (anti-GFP). Intensity measurements from ten different cells from three separate experiments were used. Intensity profiles and flagellar length measurements were measured using FIJI and the total integrated intensity was calculated by determining the total area under the curve (AUC) using custom Python scripts.

### Quantification of kinesin13mNG membrane-bound axoneme fluorescence intensity

Trophozoites expressing kinesin13mNG were prepared for live imaging as above. The segmented, spline-fit line tool in FIJI was used to trace the length of the flagellum, from the tip to the base, and measure the intensity. Measurements from at least 23 cells for each flagellar pair from two independent experiments were used. Only cells with at least two measured flagella were analyzed. Custom Python scripts were used to generate plots and statistical analyses of the data.

## Acknowledgements

This work was supported by NIH/NIAID awards 2R01AI077571-10A1 to SCD. JK is funded by the National Science Foundation grants DMR-1610737 and MRSEC-1420382, and by the Simons Foundation. SGM was supported by NIH T32 GM0007377. Plasmid pKS_mNeonGreen-N11_NEO was a gift from Alex Paredez (University of Washington, Seattle). We thank the MCB Light Microscopy Imaging Facility, a UC Davis Campus Core Research Facility, for the use of the 3i spinning disc confocal microscope (Intelligent Imaging Innovations, Inc.)and the N-SIM Structured Illumination Super-resolution Microscope (Nikon). We thank Kari Hagen for valuable editorial assistance and the Physiology course at the Marine Biological Laboratory.

---

**Supplemental Figure 1:**
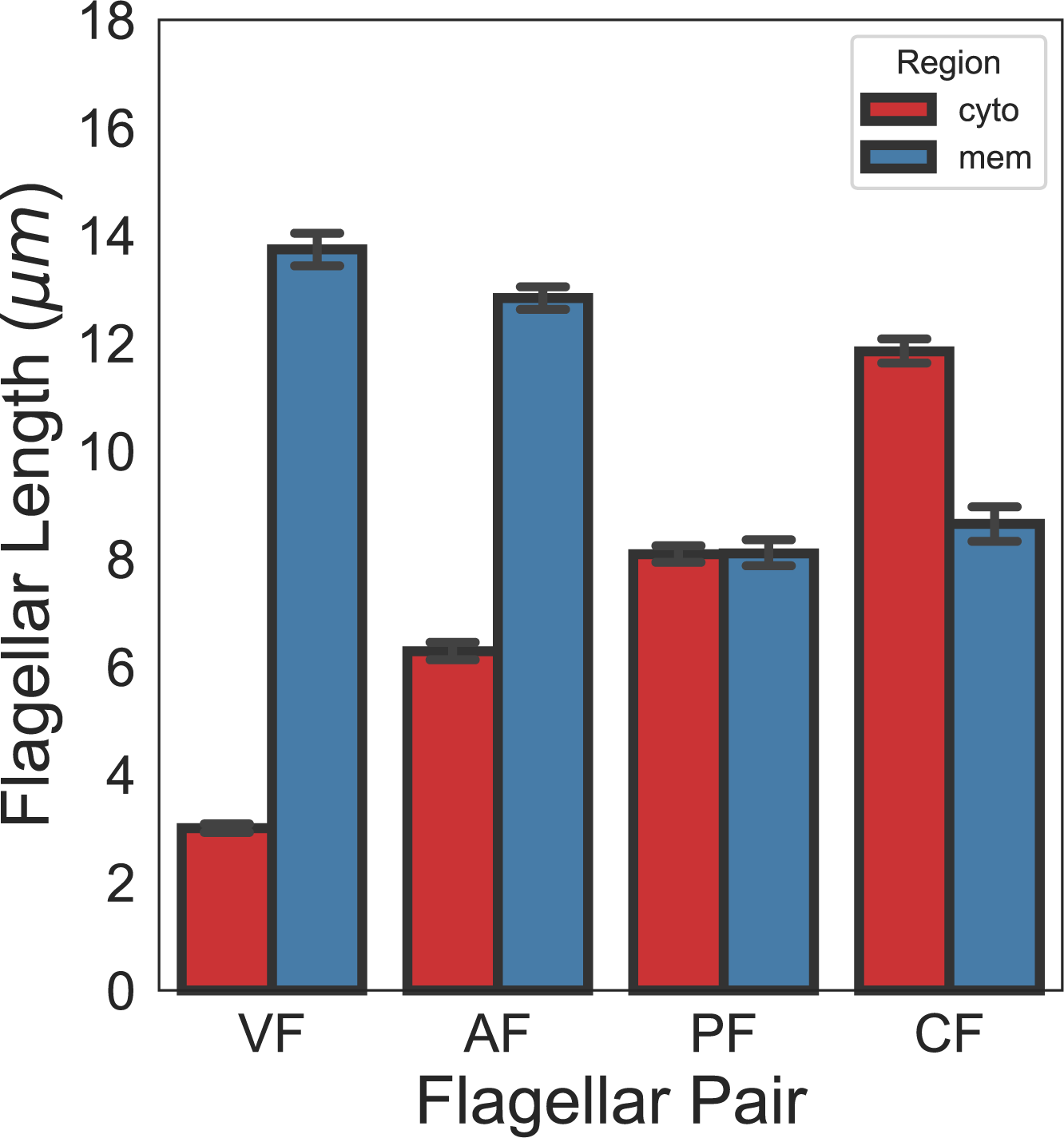
Quantification of full axoneme lengths in *Giardia lamblia*. Flagellar length quantification of membrane-bound and cytoplasmic regions of flagellar pairs of *Giardia* trophozoites expressing single-copy, integrated mNeonGreen-β-tubulin. The 95% confidence interval and average length are indicated for cytoplasmic (red) and membrane-bound (blue) regions. n ≥35 flagella for each pair from 3 independent experiments.

**Supplemental Figure 2:**
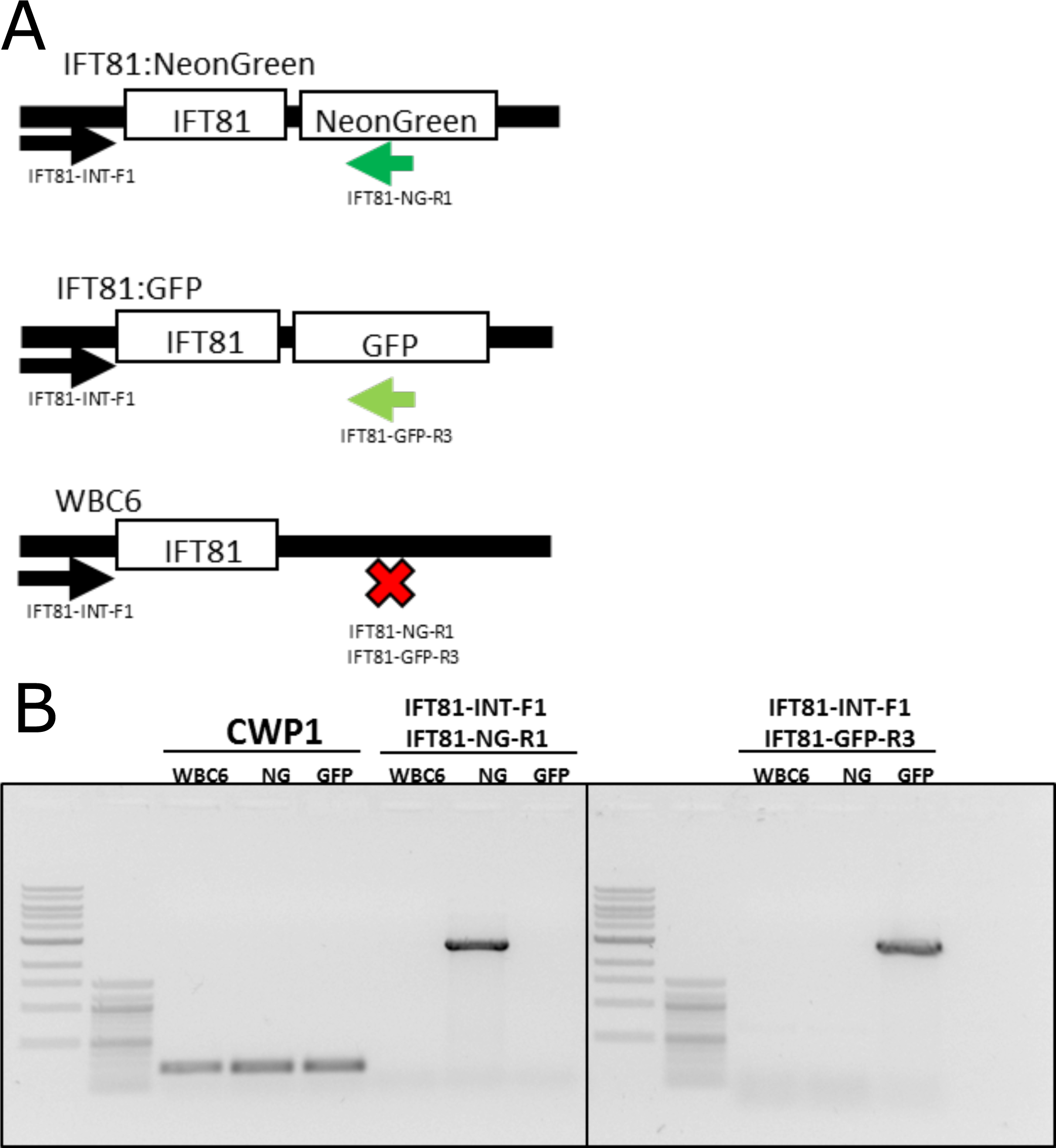
PCR validation of IFT81mNG and IFT81GFP integration into the native genomic loci. (A) Schematic representation of primers designed to detect single-copy integration of IFT81mNeonGreen and IFT81GFP constructs. (B) PCR validation of IFT81mNeonGreen and IFT81GFP integration into the native genomic locus. Specific bands are detected for each strain and positive control amplification (CWP1) is detected for all tested strains.

**Supplemental Figure 3:**
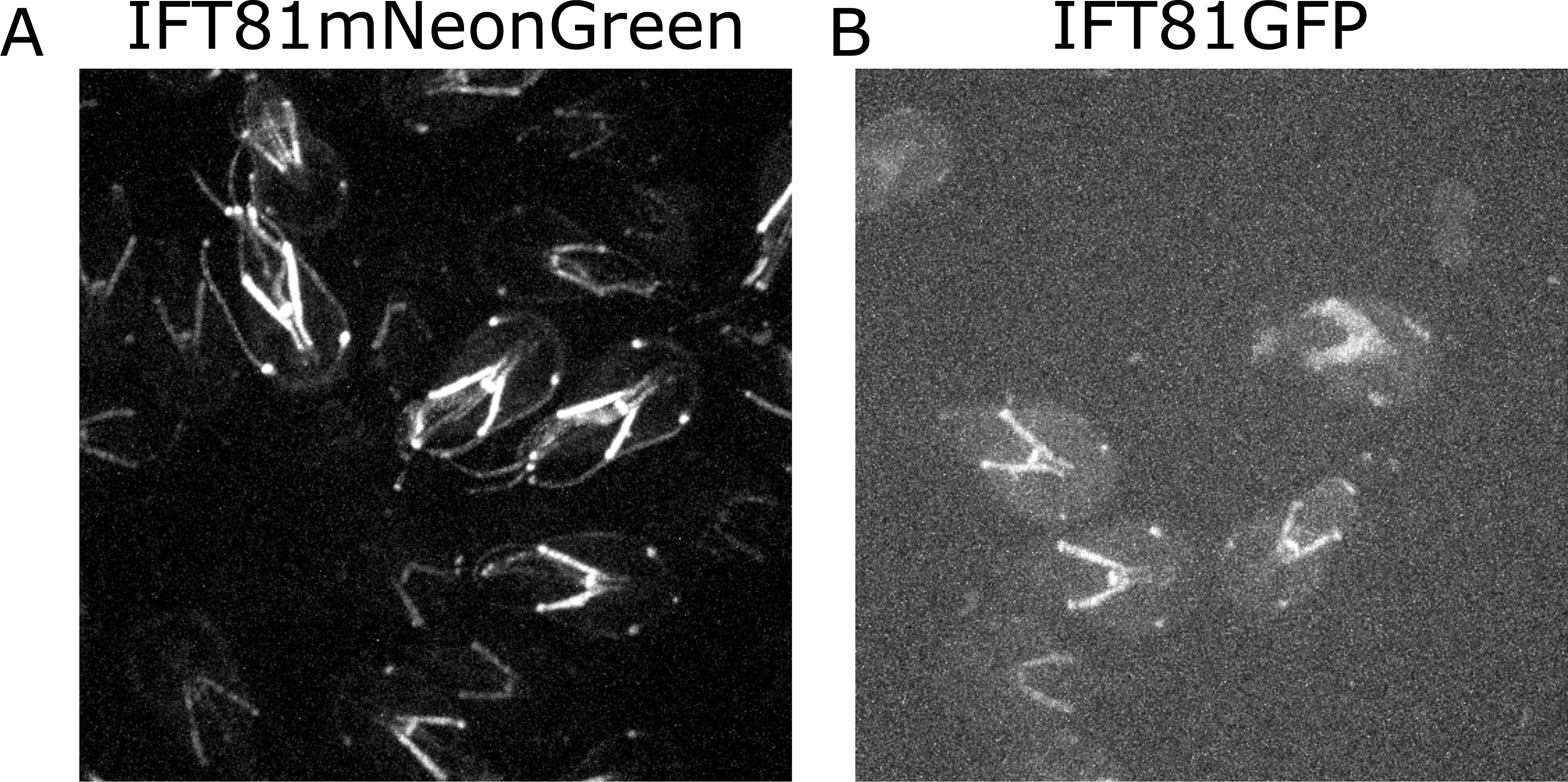
Comparison of brightness between IFT81GFP and IFT81mNG. (A) Representative image of IFT81mNeonGreen and (B) IFT81GFP. Both images are maximum intensity projections acquired with the same acquisition parameters (30ms exposure, 60% laser power).

**Supplemental Figure 4:**
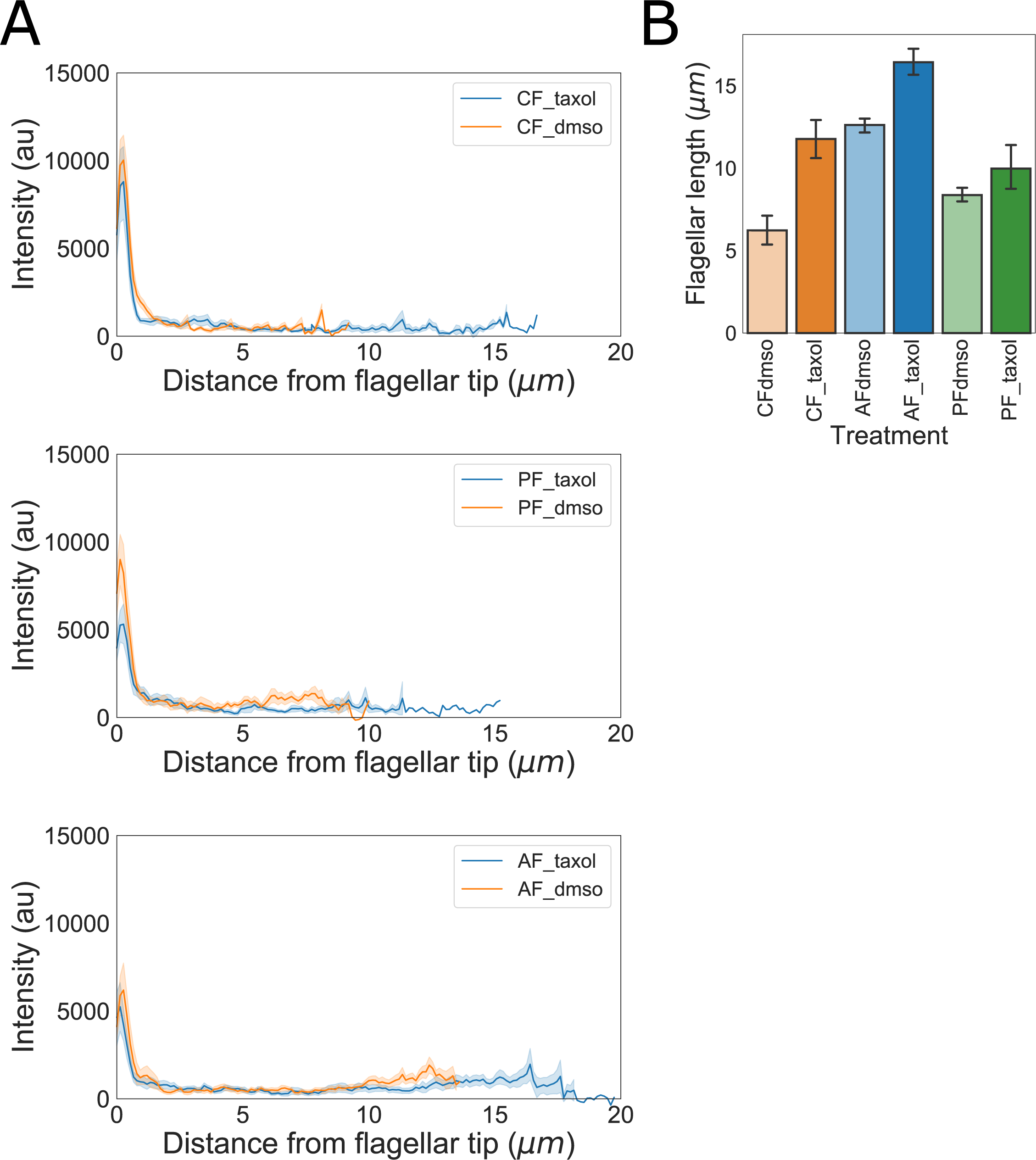
Kinesin-13mNG flagellar length changes and intensity profiles following flagellar elongation with Taxol. (A) Kinesin-13mNG intensity profiles from the flagellar tip to the base of the membrane-bound regions of caudal (top), posteriolateral (middle), and anterior (bottom) flagella. Orange traces indicate control (DMSO treated) cells and blue traces indicate Taxol treated cells. Shading indicates standard error of the mean. (B) Flagellar length of kinesin-13mNG trophozoites treated with DMSO (opaque) or 20µM Taxol (filled). n≥12 cells from two separate experiments were measured for each condition. Means and 95% confidence intervals are indicated.

Supplemental Video 1: Fluorescence recovery of IFT81mNG after photobleaching of posteriolateral cytoplasmic axonemes.

Fluorescence recovery following photobleaching of the left posteriolateral cytoplasmic axoneme in trophozoites expressing IFT81mNG. The video was recorded at 1 frame/second and is played at 10x increased speed. Time post-bleach (in minutes) is indicated in the top left corner. Scale bar, 5µm.

Supplemental Video 2: Fluorescence recovery of IFT81mNG after photobleaching of anterior and posteriolateral flagellar pores.

Fluorescence recovery following photobleaching of the right posteriolateral flagellar pore (top left) and right anterior flagellar pore (bottom right) in trophozoites expressing IFT81mNG. The video was recorded at 1 frame/second and is played at 10x increased speed. Time post-bleach (in minutes) is indicated in the top left corner. Scale bar, 5µm.

Supplemental Video 3: Tracking IFT trains in *Giardia lamblia*.

IFT train movement visualized using spinning-disc confocal microscopy in trophozoites expressing IFT81mNG. The video was recorded at ∼13 frames/second and is played in real time (indicated in the top left corner, in seconds). Scale bar, 5µm.

Supplemental Video 4: Fluorescence recovery of kinesin-13mNG after photobleaching of caudal and posteriolateral flagellar tips.

Fluorescence recovery following photobleaching of the flagellar tips of caudal (left) and posteriolateral (right) flagellar tips in trophozoites expressing kinesin-13mNG. The video was recorded at 1 frame/second and is played at 10x increased speed. Time post-bleach (in minutes) is indicated in the top left corner. Scale bar, 5µm.

