## Supplementary figures and images for "Length-dependent disassembly maintains four different flagellar lengths in *Giardia*"

### Video S1

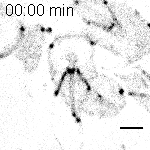

### Video S2

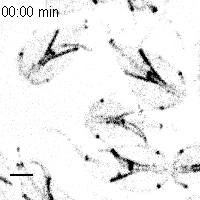

### Video S3

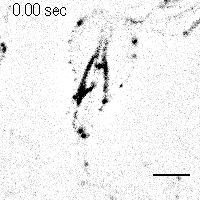

### Video S4

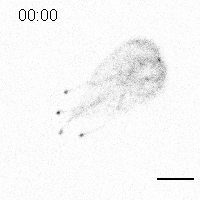
